# Tree-based differential testing using inferential uncertainty for RNA-Seq

**DOI:** 10.1101/2023.12.25.573288

**Authors:** Noor Pratap Singh, Euphy Y. Wu, Jason Fan, Michael I. Love, Rob Patro

**Affiliations:** Department of Computer Science, University of Maryland, College Park; Department of Biostatistics, University of North Carolina-Chapel Hill; Department of Genetics, University of North Carolina-Chapel Hill

## Abstract

Identifying differentially expressed transcripts poses a crucial yet challenging problem in transcriptomics. Substantial uncertainty is associated with the abundance estimates of certain transcripts which, if ignored, can lead to the exaggeration of false positives and, if included, may lead to reduced power. Given a set of RNA-Seq samples, `TreeTerminus` arranges transcripts in a hierarchical tree structure that encodes different layers of resolution for interpretation of the abundance of transcriptional groups, with uncertainty generally decreasing as one ascends the tree from the leaves. We introduce `mehenDi`, which utilizes the tree structure from `TreeTerminus` for differential testing. The nodes output by `mehenDi`, called the selected nodes are determined in a data-driven manner to maximize the signal that can be extracted from the data while controlling for the uncertainty associated with estimating the transcript abundances. The identified selected nodes can include transcripts and inner nodes, with no two nodes having an ancestor/descendant relationship. We evaluated our method on both simulated and experimental datasets, comparing its performance with other tree-based differential methods as well as with uncertainty-aware differential transcript/gene expression methods. Our method detects inner nodes that show a strong signal for differential expression, which would have been overlooked when analyzing the transcripts alone.

## 1 Introduction

RNA-Seq has become the *de facto* technology for measuring the expression profiles of different genomic features. Finding the features that differ in expression between the biological samples under different conditions, such as normal versus tumor samples, is called differential analysis and is a fundamental task downstream of quantification. The starting point of differential analysis is to accurately estimate the abundance of the features to be tested, with genes and transcripts being the most common features of interest for most RNA-Seq analysis. A gene can express multiple isoforms or transcripts due to alternative splicing, where the transcripts can comprise an overlapping set of exons and thus share sequences. Commonly used RNA-Seq quantification methods such as `RSEM` [Li and Dewey, 2011], `Salmon` [Patro et al., 2017] and `kallisto` [Bray et al., 2016] probabilistically assign each observed fragment to the transcripts using maximum likelihood or Bayesian inference methods. A probabilistic model is required because a substantial fraction of the sequenced reads can multi-map (align similarly or equally well to multiple reference transcripts) as they arise from the regions and sequences shared between the transcripts. Thus, there is ambiguity concerning the true locus of origin of such reads, leading to uncertainty when trying to quantify transcript abundances. Uncertainty can also exist for gene abundance estimates when the reads belong to shared sequences within the genes (for example, homologous genes); however, in general, gene-level estimates will be more precise compared to transcripts. One approach that estimates the uncertainty associated with the measurement of the abundance estimates is to generate additional samples called inferential replicates through different sampling strategies. These include bootstrap sampling or posterior sampling using MCMC/Gibbs sampling. Such capabilities are provided by several existing quantification methods [Li and Dewey, 2011, Bray et al., 2016, Patro et al., 2017, Turro et al., 2011].

The robustness and accuracy of any downstream analysis, such as differential testing, is directly impacted by the quality of the abundance estimates. The methods designed for differential transcript expression testing that utilize inferential replicates report a more robust performance than the methods that do not include them [Seesi et al., 2014, Mandric et al., 2017, Pimentel et al., 2017, Zhu et al., 2019, Baldoni et al., 2023]. However, if inferential replicates are incorporated, then we might observe a reduced power for the transcripts that exhibit high uncertainty since we might not be confident of the observed differential signal. In such cases, we might be able to discover more existing differential signals by aggregating transcripts that share reads together into a transcript group. This idea was first proposed in `mmcollapse` [Turro et al., 2014], also described by Roberts and Watson [Robert and Watson, 2015] and further expanded in Terminus [Sarkar et al., 2020], where these groups that contain multiple transcripts, rather than single transcripts, form the feature set for differential analysis.

Expanding upon the above motivation, we recently published `TreeTerminus` [Singh et al., 2023], which outputs a forest of transcript trees for the samples in an RNA-Seq experiment. The leaves represent the transcripts, and the inner nodes represent the aggregated group of the constituent transcripts rooted at each node. The tree encodes different layers of resolution for the interpretation of the abundance of transcriptional groups, with uncertainty generally decreasing as the tree is ascended. This tree can be utilized to find differentially expressed nodes between conditions of interest, where the nodes can consist of both leaves (transcripts) and inner nodes (transcript groups). It is possible that if the differential signal exists at a transcript, then it might also propagate at the inner nodes. A finer resolution, i.e., being as close as possible to the leaves, is, however, preferable for many analyses. The inner nodes should be only part of the output when they constitute the subsets of transcripts whose signal gets masked, primarily due to high uncertainty. Otherwise, the transcripts and not their parent nodes, should be selected. Thus, a tree-based differential testing method should be able to find such nodes in a data-driven manner.

While several tree-based differential testing methods have been proposed, they differ in the error rates they control, the underlying objective functions they seek to optimize, and the data modalities they intend to target. One class of methods focuses on testing for the global null hypothesis, meaning that for a given inner node, the null hypothesis for all the descendant leaves is true [Miecznikowski and Wang, 2023]. These methods use a top-down strategy where they descend an inner node only if the null hypothesis at that node is rejected and continue testing until the null hypothesis cannot be rejected or a leaf node is reached [Meinshausen, 2008, Goeman and Solari, 2010, Yekutieli, 2008, Lynch and Guo, 2016, Bogomolov et al., 2021]. While [Meinshausen, 2008, Goeman and Solari, 2010] control the family-wise error rate (FWER) on the rejected set, [Yekutieli, 2008, Lynch and Guo, 2016] control the false discovery rate (FDR), and [Bogomolov et al., 2021] proposes and controls a new error rate called selective false discovery rate, which is the expectation across level-specific false discovery proportion (FDP). Applying these methods can lead to accepting the null hypothesis at a node in the tree, which can consist of children that contain the differential signal but exhibit a contrasting direction of sign change between the conditions of interest (e.g. two sibling nodes for an inner node that have similar abundance and strong fold change but with opposite signs, can reduce or nullify the strength of the signal at the parent node.). This is very likely to happen when we do the hypothesis testing or compute the p-value at each node of the tree individually. We might even end up not rejecting the null hypothesis at the root node, even if there is strong evidence of the signal existing at a node situated at a lower height in the tree.

`BOUTH` [Li et al., 2022] proposes an alternative to the above approaches by defining a new hypothesis test called the modified null hypothesis and controlling an error rate called the false selection rate. It employs a bottom-up procedure and examines whether for a node the null hypothesis is true for all the descendant nodes not previously detected (detected implies that the null hypothesis is rejected) at the lower level. It identifies and outputs “driver” nodes such that none of its ancestor nodes is detected. If the driver nodes were to be analyzed, then the features might provide a coarse-grained level of analysis, especially if a clearer signal already exists at a lower height in the tree. On the other hand, it is not clear which descendant nodes corresponding to a driver node should be analyzed to provide a finer resolution of analysis.

Another category of methods aims to leverage the tree structure to enhance power and control the FDR at the leaf level [Xiao et al., 2017, Huang et al., 2021, Bichat et al., 2022]. `StructFDR` [Xiao et al., 2017]) and zazou [Bichat et al., 2022] initially convert the p-values into z-scores and then propose smoothing on the leaves using a tree-based correlation structure. `treeclimbR` [Huang et al., 2021] proposes multiple candidate sets, performs multiplicity correction on each set, and selects a final candidate set based on a set of criteria. It rejects the null hypothesis for all the leaves belonging to the inner nodes in the selected candidate set. TEAM [Pura et al., 2019] also proposes another approach for controlling FDR on an aggregated tree but in a different context compared to the methods described so far. The leaves in the tree are called bins, which consist of cells from both conditions. The tree structure aggregates bins and not the features as we go higher up the tree. The null hypothesis tested at each bin is if the PDF of features is different between the two populations in that bin. If the null hypothesis at a bin can’t be rejected at a lower level, then such bins are aggregated and the hypothesis is tested at the aggregated bin. Importantly, if the aggregated bin is rejected at a higher level in the tree, then the method rejects all the underlying leaf bins. Thus, all these methods cannot be directly applied to our use case. There is uncertainty in measuring the abundance estimate of the transcripts, thus we do not know which transcript corresponding to the node may be differentially expressed. As a result, passing the information from an inner node to all the descendant leaves might lead to an exaggeration of the false positives or missing the branches where there exists a signal at a node located at a higher height in the tree, to control for the false positives on the leaf set.

There exists another class of methods that fit regression models to compositional data for the covariates using a tree-guided regularization in a maximum likelihood (`trac` - [Bien et al., 2021]) or Bayesian setting (`tascCODA` - [Ostner et al., 2021]). For `trac`, the parsimony of feature selection (where more nodes at greater heights result in a lesser number of total nodes) depends on the weight assigned to regularization during model fitting. For both `trac` and `tascCODA`, when an inner node is selected, its learned coefficients are passed equally to its underlying leaves. In addition to some of the limitations mentioned above that also exist for these methods, both these methods do not produce p-values. This can complicate comparisons with other methods. `trac` requires cross-validation to determine appropriate hyperparameters for the model, thus requiring a substantial number of samples, which may not be feasible for many RNA-Seq experiments. `tascCODA` also does not scale computationally for datasets with a large number of features, such as RNA-Seq.

To overcome some of the above limitations, we introduce our differential testing method, mehenDi, designed to output a set of nodes called the selected nodes from the tree. The selected nodes are differentially expressed and can comprise both transcripts and inner nodes, with no two nodes having an ancestor/descendant relationship. It utilizes the tree/s obtained from `TreeTerminus`, and applies the Swish method for hypothesis testing, accounting for inferential uncertainty (though it is conceptually compatible with other differential testing methods that incorporate inferential uncertainty). The selected nodes are determined in a data-driven fashion by traversing the tree in a top-down manner using a set of rules to maximize the signal that can be extracted from the data while controlling for the uncertainty associated with measuring the abundance estimates. The selected inner node represents a subset of transcripts whose signals are masked at lower levels due to inferential uncertainty, thus we do not report the differential status of the constituent transcripts. We applied our method to experimental and simulated datasets, comparing its performance with the other tree-based differential testing methods and uncertainty-aware differential transcript/gene estimation methods. Our method demonstrates improved sensitivity compared to alternative approaches, particularly when the features consist only of transcripts or transcript groups. It identifies inner nodes exhibiting a strong signal for differential expression between the conditions, which might be overlooked when examining their underlying transcripts alone. The differential signal can often be restricted to only a subset of the annotated isoforms of a gene. Thus, our approach also mitigates the potential over-aggregation of signals if reporting is done at the gene level. The R package implementing mehenDi is freely available and open-source (under the MIT license) and can be obtained from https://github.com/NPSDC/mehenDi.

## 2 Results

### 2.1 Overview

There can exist substantial uncertainty in estimating the abundance of the transcripts due to the presence of reads that can map equally well to shared regions of sequence between transcripts. This quantification uncertainty, in turn, can impact the quality and robustness of differential analysis using transcripts as features. mehenDi provides a different approach to tackle this problem for RNA-Seq data. For a given set of samples belonging to an RNA-Seq experiment, `TreeTerminus` (Singh et al. [2023]) arranges transcripts in a hierarchical tree structure, based on quantification uncertainty (Figure 1A). mehenDi has been designed to perform differential testing directly over the tree/s produced by `TreeTerminus`.

**Figure 1:**
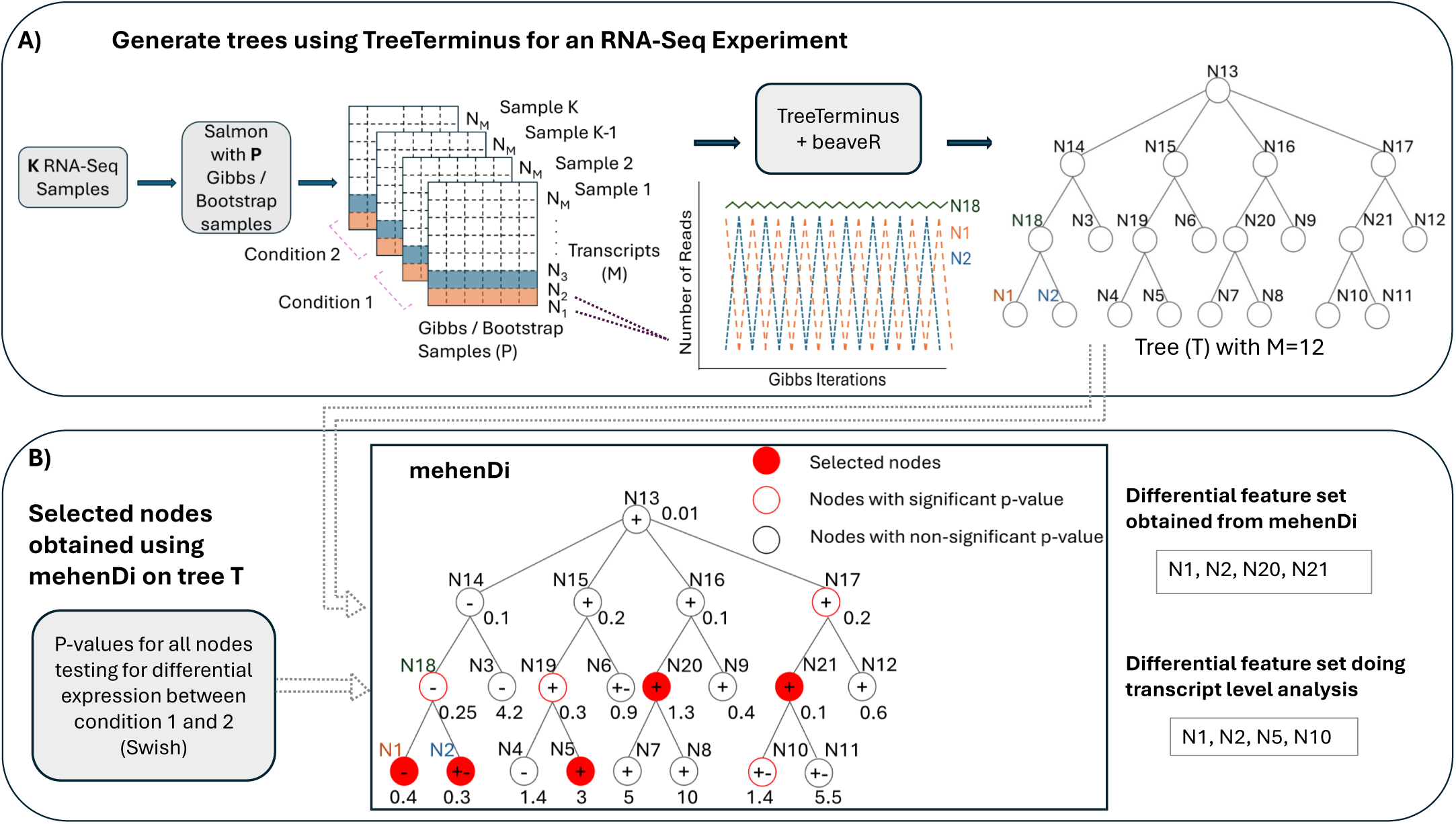
Schematic overview of `mehenDi`. A) Given a set of RNA-Seq samples, `TreeTerminus` creates a tree that encodes the uncertainty structure. B) Given a tree, `mehenDi` outputs a set of selected nodes. The selected are differentially expressed between the conditions of interest and can consist of transcripts and inner nodes, with n. Using an example, with a tree having 12 leaves to demonstrate `mehenDi`. A highlighted node implies that it is selected by `mehenDi`, with a red outlined node indicating that the node has a significant p-value. The +, − on a node indicates the direction of sign change between the conditions, whereas +− indicates we are unsure about the direction. The values plotted along the nodes represent their mean inferential relative variance across the samples (meanInfRV). The meanInfRV threshold in this example is 0.4. The nodes N1 and N2 are chosen instead of node N18 since the meanInfRV of one of its children, node N2, is below the threshold of 0.4. The node N5 is chosen instead of the node N19 since the children have opposite sign changes. Node N20 is the only significant node in its branch and meets all the required criteria. Node N21 is picked since it has at least one non-significant child node.

As input, mehenDi requires the p-values (computed through an uncertainty-aware differential testing method), the direction of sign change and mean inferential relative variance (metric to quantify uncertainty) for all nodes in the tree (Figure 1B). It uses a top-down procedure to output a set of selected nodes that are differentially expressed between the conditions of interest, which seek to maximize the signal that can be extracted from the data while controlling for the uncertainty associated with estimating the transcript abundances. The selected nodes can include transcripts and inner nodes (transcript groups), with no two nodes having an ancestor/descendant relationship. Intuitively, mehenDi allows reporting differential expression at the level of transcript groups (groups of structurally related transcripts) and finds signals that may have been too weak to determine at the level of individual transcripts and which at the gene level, may involve the aggregation of other structurally dissimilar transcripts not involved in the differential regulation. Figure 1B also provides a toy example demonstrating mehenDi. The exact procedure is described in detail in the Methods section.

### 2.2 Running mehenDi on null simulation

We recommend computing p-values for leaves and inner nodes separately (see Methods) before running mehenDi. Through this analysis, we want to demonstrate why we recommend computing the p-values in the specified manner. We also aim to evaluate the false positive rate for mehenDi specifically comparing it to transcript-level differential analysis.

The simulation framework involves simulating reads from the human transcriptome for 12 samples belonging to two groups (which represent the conditions of interest), with 6 samples in each group. In the null simulation experiment, the fold change between the groups is fixed at 1 for all transcripts. We create a total of 10 null simulation experiments. The framework has been described in detail in the Methods section. Thus, by design, the simulation has no true signal for differential expression for the transcripts and any node deemed as significant by a method would be a false positive.

#### 2.2.1 Inner nodes have smaller p-values if the hypotheses testing is carried out on all the nodes of the tree simultaneously

For each null simulation, we ran Swish on all the tree nodes simultaneously, and then the computed p-values across all the null simulations were concatenated into one vector. As expected, the distribution of the p-values across all the nodes is uniform (Figure 2A i). However, if we look at the distribution of the p-values at the leaves, there is a shift in the distribution towards the right for the leaf p-values and the left for the inner node p-values (Figure 2A ii,iii). Ideally, irrespective of the subset of features that are picked, the distribution should be uniform. This happens because the width of the distribution of the test-statistic which is used by Swish is wider for the inner nodes compared to the leaves (Figure 2A iv). Thus, if the p-values are generated together for all nodes, we might inflate false positives for the inner nodes and false negatives for the leaves. We thus compute the p-values for the leaves and inner nodes separately and, as expected, both of these show a uniform distribution (Supplementary figure S1).

**Figure 2:**
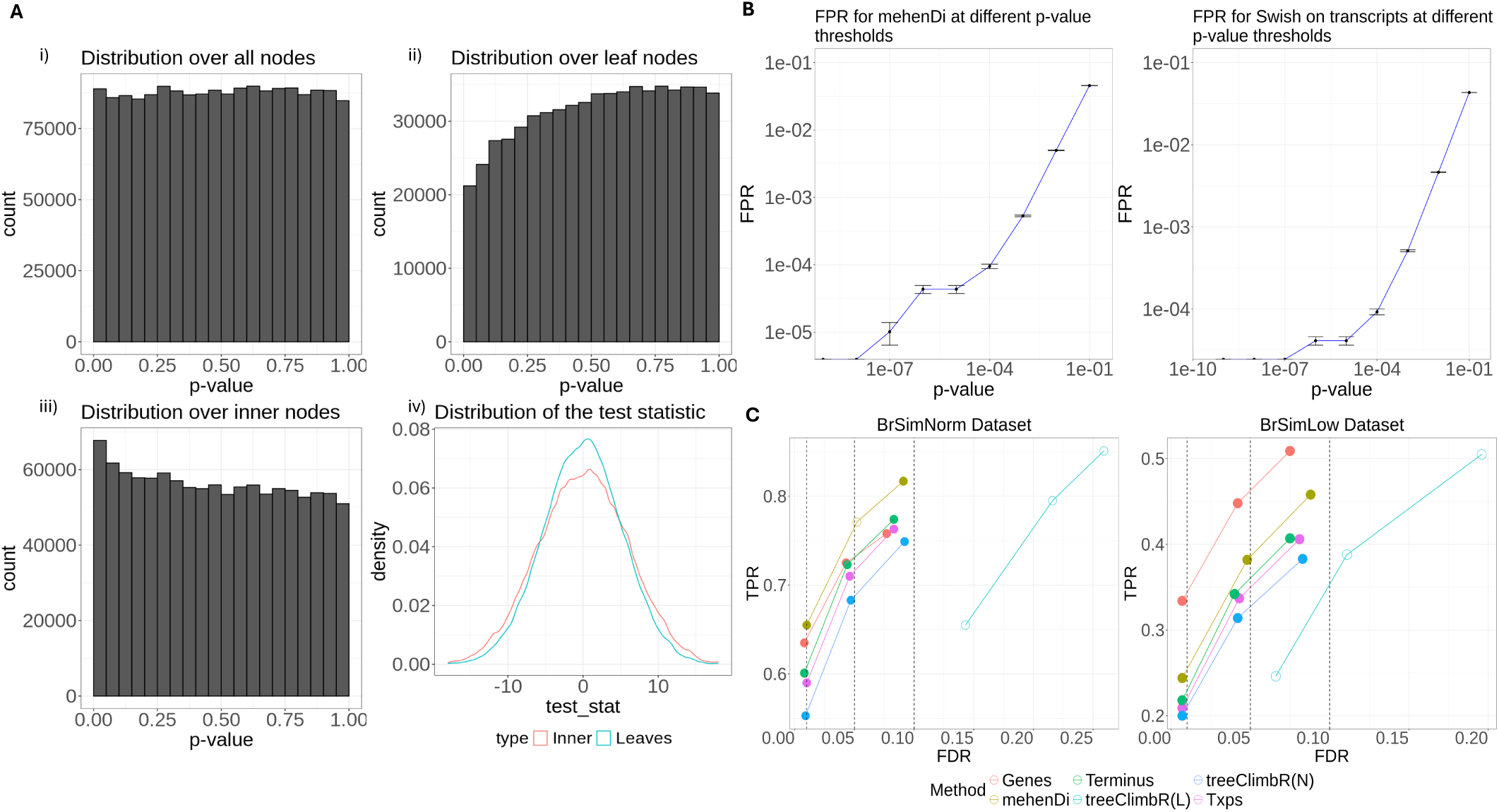
A) Distribution of p-values and test-statistic for Swish for the null brain simulation. i) Distribution of the p-values on the entire node set over the tree when all hypotheses are tested together using Swish. ii) Distribution of the p-values over the leaf nodes. iii) Distribution of the p-values over the inner nodes. iv) Distribution of the test-statistic on the inner and leaf nodes, which is used by Swish to generate the p-values. B) False Positive Rate (FPR) obtained for the null simulations by varying the p-value threshold. The bar along the y-axis represents the variation observed across the 10 simulations. C) True Positive Rates and Empirical False Discovery Rates at the different nominal FDR thresholds across the different methods and entities for the BrSimNorm and BrSimLow Dataset. Both the metrics have been rounded to 3 decimal places.

#### 2.2.2 The False Positive Rate (FPR) obtained by mehenDi on the tree is comparable to running Swish on only the transcripts

We next looked at the False Positive Rate (FPR) reported by mehenDi on the null simulations and compared it with the FPR obtained by running Swish only on the transcripts. We varied the p-value threshold from 1*e* − 9 to 1*e* − 1, increasing the threshold by the power of 10 and then computed the FPR across the 10 simulations, shown in Figure 2B. The empirical FPR that is obtained across both settings is comparable to each other. This experiment demonstrates that mehenDi should not exaggerate the false positive rate compared to Swish on transcripts.

### 2.3 Comparing the methods on simulated datasets with signal

We next benchmark and analyze the simulated datasets with differential signals i.e. `BrSimNorm` and `BrSimLow` datasets (see Methods), which again consist of 12 samples belonging to two groups. Specifically, transcripts in `BrSimLow` show a relatively weaker signal for differential expression on average compared to `BrSimNorm`.

#### 2.3.1 Features obtained from mehenDi show good sensitivity while controlling for FDR

We compare the features obtained after differential testing using different methods. The features consist of genes (Genes), transcripts (Txps, `treeclimbR`(L)) and transcript-groups (Terminus, mehenDi, `treeclimbR`(N), BOUTH). Txps refers to the transcripts obtained by running vanilla Swish only on the transcripts. When `treeclimbR` selects an inner node as a candidate node, this implies that all of its descendant transcripts are also differentially expressed. Thus, in our analysis, we compare two feature sets corresponding to `treeclimbR`: `treeclimbR`(N), which consists of only the best candidate nodes, and `treeclimbR`(L), which consists of the underlying transcripts belonging to the nodes. This enables us to compare the performance of `treeclimbR` at both the node and transcript levels. On all the simulated and experimental datasets, BOUTH outputs the root node as the driver. Since the root node represents the aggregation of all the transcripts, interpreting the results from BOUTH does not provide any information. We first look at the number of true positive transcripts that are covered by the different methods on the simulated datasets (Supplementary figure S2, S3) (excluding BOUTH). We observe that mehenDi yields the largest number of true positives, followed by `treeclimbR`.

We next compute TPR and FDR at different nominal FDR thresholds on the feature sets obtained from the different methods (Figure 2C). For the `BrSimNorm` dataset, `treeclimbR`(L) and mehenDi have the largest TPR, while for the `BrSimLow` dataset, Genes has the highest TPR, followed by `treeclimbR`(L) and mehenDi. `treeclimbR`(L) has an inflated FDR on both datasets (since all transcripts belonging to a candidate node might not be true positives), whereas `treeclimbR`(N), while controlling for FDR, does not seem to lead to that much overall improvement in sensitivity compared to Txps. mehenDi seems to have a good sensitivity while controlling for the FDR at the same time, with FDR only going slightly beyond the nominal FDR threshold of 0.05 for the `BrSimNorm` dataset. We have excluded the results of BOUTH in the figure since it outputs the root node at all the different nominal FDR values we have tried in our experiments. By design, the root node is not significant in the simulation and thus the TPR and FDR for BOUTH are 0 and 1 respectively.

We also individually varied the parameters `minP` and `mIrvThresh` (see Methods) for mehenDi and observed how the performance changed for the `BrSimNorm` dataset, shown in Supplementary figure S4. We first kept the parameter `mIrvThresh` constant to its default value 0.4 and varied `minP` from 0.6 to 0.9, incrementing it at an interval of 0.05. We however don’t observe much change in the performance, with a very slight increase in TPR and FDR as `minP` is increased. We next kept the parameter `minP` constant to its default value 0.7 and varied `mIrvThresh` from 0 to 1, incrementing it at an interval of 0.1. The FDR and TPR both decrease when `mIrvThresh` is increased from 0 to 1, with the FDR for 0.01 controlled after `mIrvThresh` 0.3, for 0.05 after `mIrvThresh` 0.6 and for 0.1 after `mIrvThresh` 0.2. At the lower values of `mIrvThresh`, mehenDi becomes less conservative and prefers nodes with larger heights, which increases the FDR. Higher height nodes are also preferred for larger values of `minP`, since the nodes downstream are more likely not to be assigned a confident sign change. Further, more nodes located at different branches in the tree are also observed for lower values of `mIrvThresh` and higher values of `minP`, compared to selected nodes obtained at the default parameters.

It is important to note that since the underlying features across the methods are different, these metrics may not be directly comparable. However, we obtain some information about their respective performances. We next investigate the increased sensitivity shown by mehenDi, by doing a pairwise comparison with the other methods on the `BrSimNorm` dataset.

#### 2.3.2 Features output by mehenDi covers more true positive transcripts compared to Terminus

Next, we perform a pairwise comparison between the features output by mehenDi to Terminus obtained at 0.01 nominal FDR. Specifically, we examine the unique true positive transcripts that have been covered by the output features in comparison to one another. We find that features output by Terminus candidate nodes correspond to only 31 unique true positive transcripts, which map to 26 Terminus groups. For mehenDi, transcripts descending from its selected nodes yield a total of 681 unique true positive transcripts and correspond to 473 nodes. While the general expectation might be that most of these additional selected nodes in mehenDi would be as a result of more aggregation, we find 27 nodes for which the signal exists at a lower level in mehenDi that get lost in Terminus due to overaggregation. We provide some examples of such cases in Figure 3A, Supplementary figure S5 and S6. Similarly, there are 103 such groups (inner nodes in the tree) in mehenDi that contain at least one underlying descendant transcript, which was differentially expressed. The differential signal is lost for these transcripts in Terminus, either because the transcripts were not grouped at all or not aggregated enough to produce the signal. In Figure 3B, Supplementary figure S7 and S8, we show some examples of such cases.

**Figure 3:**
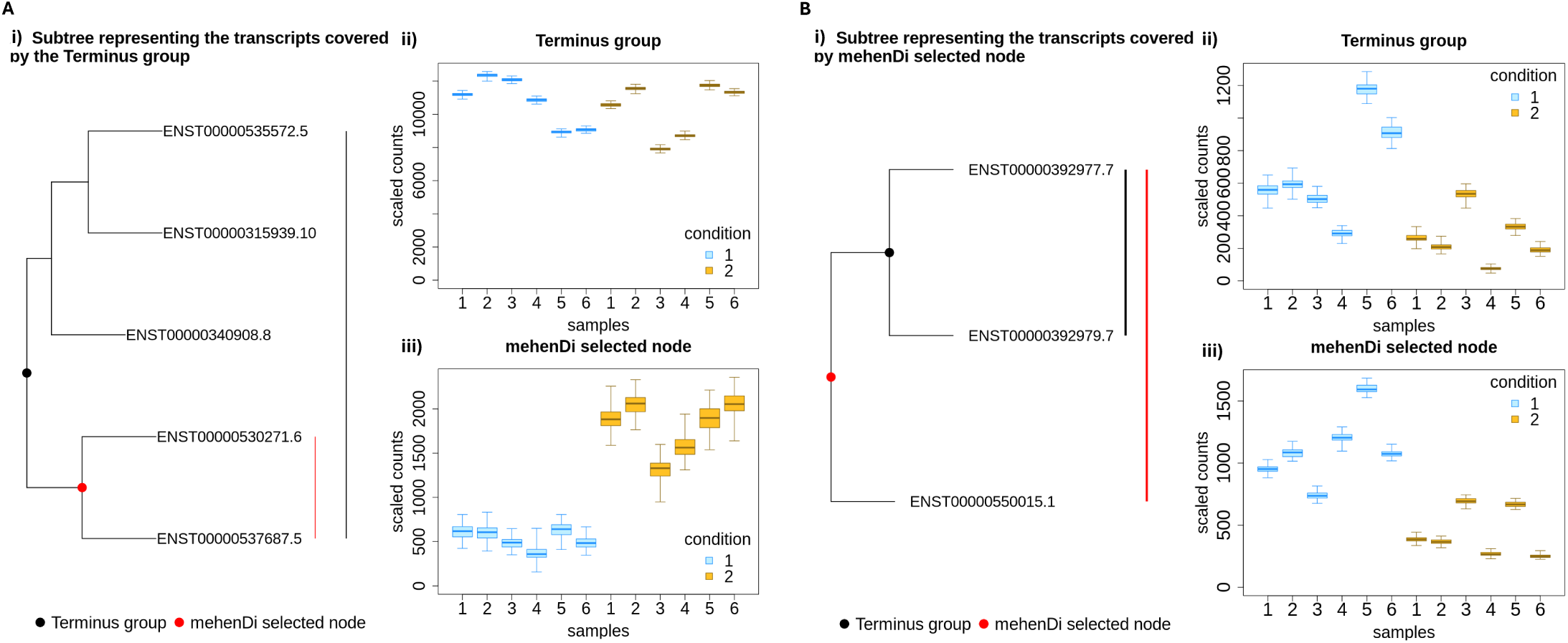
A) Example of a `mehenDi` node that is overaggregated in Terminus for the `BrSimNorm` dataset. i) Subtree representing the transcripts covered by the Terminus group. ii) Inferential replicates for the Terminus group. iii) Inferential replicates for the selected node output by `mehenDi`. B) Example of a `mehenDi` node that is not aggregated enough in Terminus for the `BrSimNorm` dataset. i) Subtree representing the transcripts covered by the `mehenDi` group. ii) Inferential replicates for the Terminus group. iii) Inferential replicates for the selected node output by `mehenDi`.

#### 2.3.3 mehenDi finds more unique branches in the tree compared to treeclimbR

We next perform a pairwise comparison between `treeclimbR` and mehenDi at the 0.01 nominal FDR on the `BrSimNorm` dataset. We first inspect the relative height of the nodes in Supplementary table S1. We observe more higher-level nodes for `treeclimbR` compared to mehenDi, with the maximum height of the output node being 19 and 14 in `treeclimbR` and mehenDi, respectively. It is not trivial to compare these two methods when the nodes output by them are located at different heights and when the differential signal can exist at multiple levels within the same branch in the tree. We thus try to identify the branches unique to a method using the approach below. For example, imagine that we want to find the nodes unique to mehenDi. To do so, we first extract the nodes that are output by both of these methods (`treeclimbR`, mehenDi). From the remaining selected nodes (excluding common nodes) that are output by mehenDi, we remove all nodes *i* for which there exists a node *j* that is an output of `treeclimbR` and is either an ancestor or descendant of node *i* in the tree. The selected nodes that are left represent the unique branches of mehenDi. We repeat a similar process for `treeclimbR`, to find the unique branches of `treeclimbR`. We find 199 such nodes unique to mehenDi that map to 258 true positive transcripts, while for `treeclimbR` we find only 2 such nodes that map to 4 true positive transcripts. Thus mehenDi finds more unique branches in the tree which in turn can enable us to recover the signal from more true positive transcripts.

#### 2.3.4 FDR is not controlled across all the methods on the unique nodes

Since each method can output unique nodes, we next assess the error rate only on the unique branches/nodes by comparing the methods pairwise. The unique nodes are extracted using the approach described above for the different nominal FDR thresholds. Specifically, we extract the nodes for pairwise comparison between mehenDi versus `treeclimbR`, and `treeclimbR` versus Txps. We provide the observed FDR and the total number of unique nodes for these comparisons in Supplementary table S2, S3. The multiple hypothesis correction methods aim to control the average false discovery proportion of the overall set of rejected hypotheses. However, the error guarantee might not necessarily hold on a specific subset, which in this case are the unique nodes corresponding to each method. We observe an inflated observed FDR for the unique features corresponding to Txps and mehenDi, although it decreases for lower nominal FDR thresholds. The observed FDR is also inflated for `treeclimbR` on pairwise comparison with Txps, containing significantly fewer unique nodes. We next try to see if we can reduce the FDR by filtering the unique set based on the threshold of absolute log fold change between the conditions of interest. We plot this in Figure 4, Supplementary figure S9 and S10. We see that the observed FDR starts decreasing as the effect size increases and then again starts to increase. The FDR again starts increasing because, after a certain threshold of effect size, only a few nodes are left after filtering. As a result, the false positive rate increases more dramatically due to a small denominator, leading to a heavier penalty for each incorrect prediction. We thus recommend using a conservative nominal FDR threshold with an appropriate effect size when the novel features unique to any method are analyzed.

**Figure 4:**
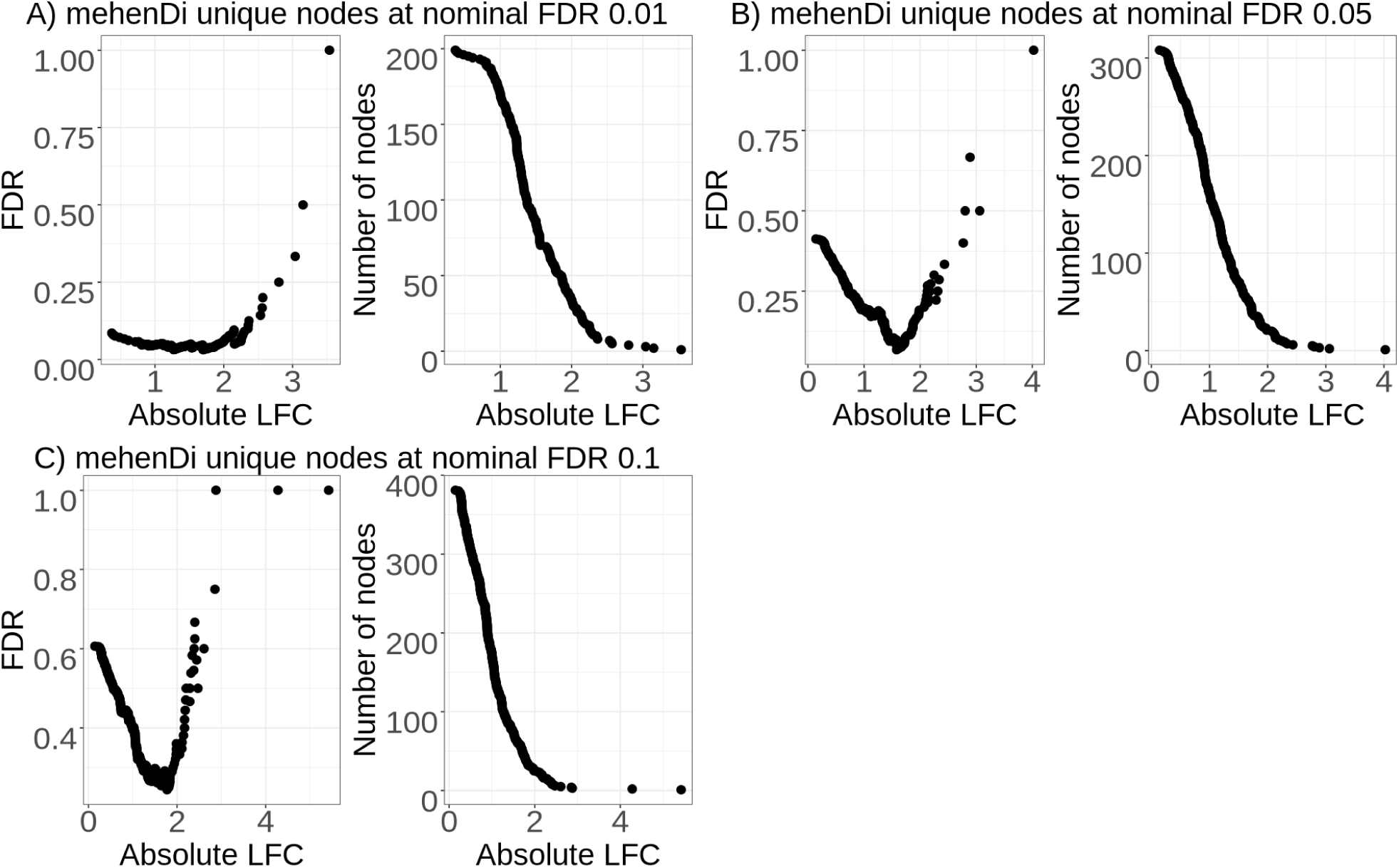
Examination of error metrics for the unique nodes obtained for `mehenDi` when doing `mehenDi` vs `treeclimbR` analysis at the different nominal FDR thresholds for the `BrSimNorm` dataset. We vary the magnitude of log fold change (LFC) and plot the empirical FDR and the total number of nodes that are left after filtering the unique nodes based on LFC.

### 2.4 Analyzing the MouseMuscle dataset

We next analyzed the `MouseMuscle` dataset which consists of 6 samples each from EDL and MAST muscle tissues (see Methods). The first two dimensions of the PCA using the top 1000 variable genes for the dataset are shown in Supplementary figure S11.

#### 2.4.1 Increase in variation in the parameter values compared to the default setting leads to larger distances between the selected nodes

We first varied the parameters of mehenDi individually and computed the distance of the selected nodes obtained at those parameters from the tree with the selected nodes obtained at the default parameters (Supplementary Figure S12). For both `minP` and `mIrvThresh`, when the parameter values are closer to the threshold, the distance is lower and increases as the parameters are varied in both directions. The distance increases more sharply when `mIrvThresh` is varied compared to when `minP` is varied. As seen in the `BrSimNorm` dataset, we also observed nodes belonging to distinct branches of the tree and located at higher heights, at low values of `mIrvThresh` and high values of `minP`. The method for computing the distances between the nodes has been described in Supplementary section S1.3

#### 2.4.2 mehenDi can find nodes for genes for which the differential signal could not be detected at the transcript-level

We next compared the nodes obtained by mehenDi at the default parameter values with the different features and the nodes obtained across the other methods. The total number of differentially expressed genes (DEGs), differentially expressed transcripts (DETs) at the different nominal FDRs and the total number of transcripts that are covered by the nodes/groups output by the different methods at the 0.01 nominal FDR are shown in Supplementary table S5, S6 and Supplementary figure S13. We observe that mehenDi-reported nodes cover more transcripts compared to `treeclimbR`-reported nodes. The distribution of the node heights output by mehenDi is shown in Supplementary table S7, whereas the distribution of the number of unique genes that the transcripts output by mehenDi map to is provided in Supplementary table S8 for the different nominal FDR thresholds. We observe that the majority of the output nodes consist of transcripts. Further, most of the transcripts covered by these nodes map to only one gene. We find many genes that are called significant when differential testing is carried out on the genes, but the underlying transcripts are not called as significant when the testing is performed on the transcripts (Supplementary table S9). Similarly, when differential testing is carried out on the transcripts, some of the significant transcripts map to non-significant genes (Supplementary table S10). Some of the selected nodes output by mehenDi neither map to a significant gene nor contain a single underlying transcript that is significant (Supplementary table S11). We also find many selected nodes in mehenDi that map to significant genes, but the genes do not contain a single transcript that is called significant during transcript-level differential analysis (Table 1). We provide some examples for these genes in (Figure 5, Supplementary figure S14, S15, S16, S17 and S18) obtained for 0.01 nominal FDR. For example, we can see that for the gene *Prrg1*, the transcripts highlighted in red (which underlie the selected node) constitute a very similar set of exons but vary in length. This is reflected in the tree structure in Figure 5A ii, where the transcripts `ENSMUST000000114024.8` and `ENSMUST000000177904.7` are aggregated first, followed by transcript `ENSMUST000000114025.7`. Thus, in Figure 5A iii we see that the transcript `ENSMUST000000177904.7` has a high inferential relative variance (`infRV`). This also affects the quality of the signal, which gets stronger at the selected node level constituting the aggregated transcripts (Figure 5A iv).

**Figure 5:**
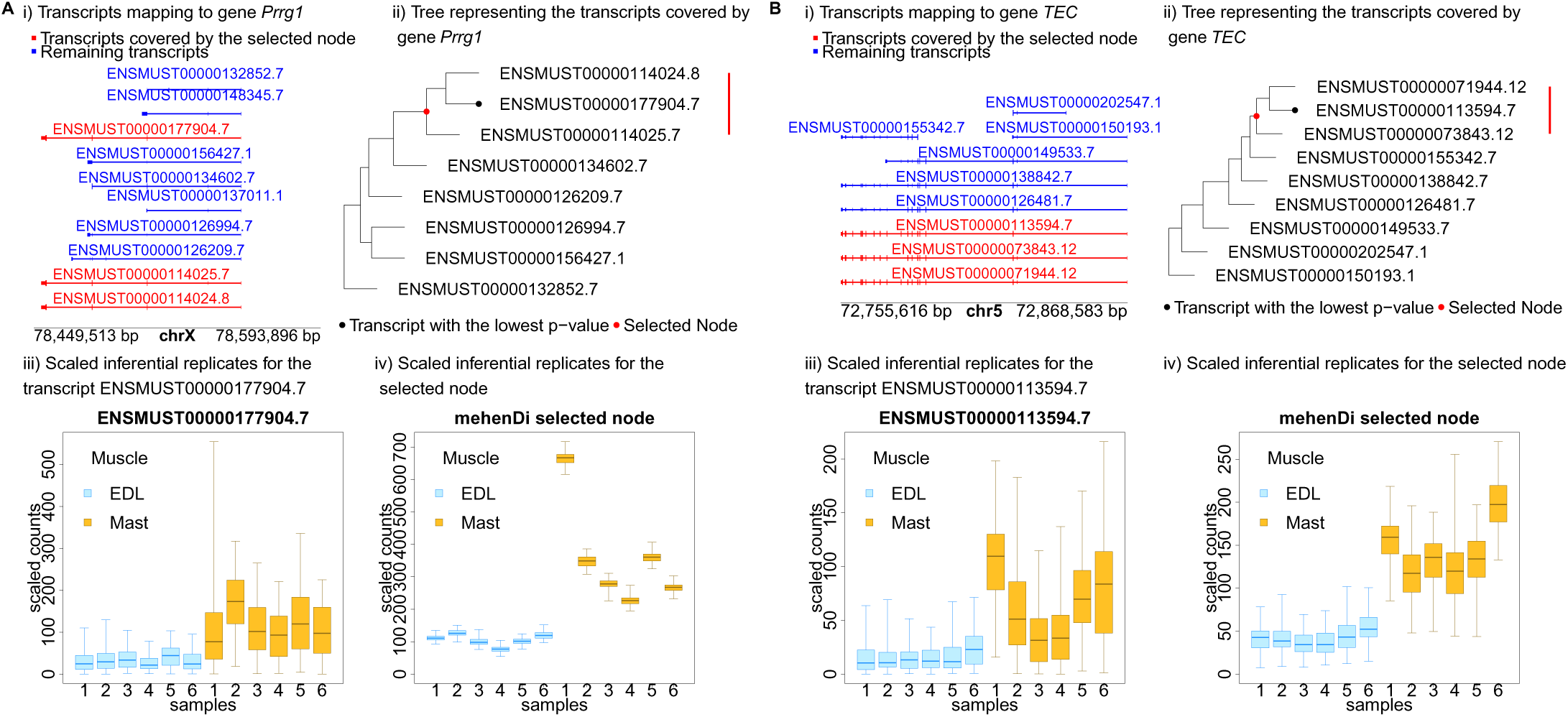
Examining the transcript profile for the gene *Prrg1* (A) and *Tec*(B) in the `MouseMuscle` dataset. i) Transcripts in a pileup style. ii) Tree representing the transcripts covered by the gene, with the red node representing the transcripts covered by the mehenDi selected node. iii) Inferential replicates for the transcript which had the lowest p-value among all the transcripts in the tree. iv) Inferential replicates for the mehenDi selected node.

**Table 1:** Number of mehenDi nodes that map to a significant gene but the gene doesn’t consist of any significant transcripts for the MouseMuscle and the ChimpBrain dataset respectively.

| Nominal FDR | Number of *mehenDi* Nodes |  |
| --- | --- | --- |
|  | *MouseMuscle* | *ChimpBrain* |
| 0.01 | 309 | 407 |
| 0.05 | 428 | 194 |
| 0.10 | 465 | 213 |

### 2.5 Analyzing the ChimpBrain dataset

We also assessed the performance of mehenDi on the ChimpBrain dataset, which contains 5 samples each from both cerebellum and medial dorsal nucleus of thalamus (see Methods). The number of genes and transcripts that are differentially expressed between the conditions at the different FDR thresholds is provided in Supplementary table S12 and S13 with the total number of features across the different methods shown in Supplementary figure S19. We again observe that the mehenDi-reported nodes cover the largest number of transcripts. The distribution of the node heights output by mehenDi is shown in Supplementary table S14, whereas the distribution of the number of unique genes that the transcripts output by mehenDi map to is provided in Supplementary table S15 for the different nominal FDR thresholds. We observe that the majority of the output nodes consist of transcripts. Further, most of the transcripts covered by these nodes map to only one gene.

#### 2.5.1 mehenDi can find nodes for genes for which the differential signal could not be detected at the transcript-level

As observed for the `MouseMuscle` dataset, we have many examples for both cases where significantly called transcripts do not necessarily map to significantly called genes and vice versa for significant called genes (Supplementary table S16, S17). Similarly, we find some of the nodes reported by mehenDi map to non-significant genes and also don’t contain a single transcript that is called significant (Supplementary table S18).

On the other hand, there also exist many other mehenDi nodes that map to those significant genes whose underlying transcripts were not called significant (Table 1). We report some of these cases in Figure 6, Supplementary figure S20, S21, S22, S23, S24 and S25 for the 0.01 nominal FDR. For example, for gene *EYA4* in Figure 6B, we observe that the transcripts ENSPTRT00000034383 and ENSPTRT00000104666 differ by only 1 exon, and the selected group that aggregates both these transcripts has lower uncertainty and shows an increased signal, compared to the transcript ENSPTRT00000104666 which has high uncertainty.

**Figure 6:**
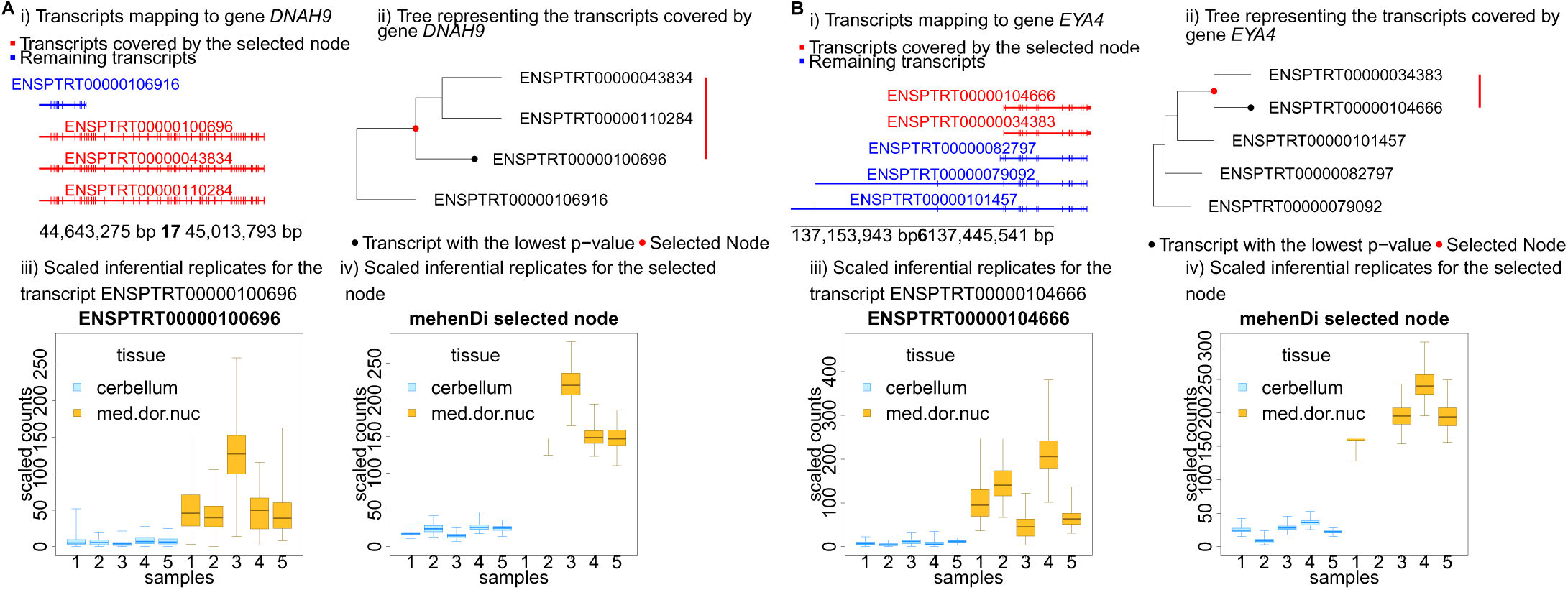
Examining the transcript profile for the gene *DNAH9* and *EYA4* in the ChimpBrain dataset. i) Transcripts in a pileup style. ii) Tree representing the transcripts covered by the gene, with the red node representing the transcripts covered by the mehenDi selected node. iii) Inferential replicates for the transcript ENSMUST00000104666, which had the lowest p-value among all the transcripts in the tree. iv) Inferential replicates for the mehenDi selected node.

### 2.6 Comparison of running time

We compared the runtime of `treeclimbR` and mehenDi on the `BrSimNorm` and the `MouseMuscle` datasets. For `treeclimbR`, two functions, namely getCand and evalCand, have to be called to obtain candidates. We executed the getCand function once and then ran the evalCand function at the different nominal FDR thresholds. mehenDi is parallelizable and supports multiple cores. In our comparison, as shown in Table 2, we presented the run time for getCand computed once and the runtime for evalCand on each nominal FDR threshold. Additionally, we showcased the time taken by mehenDi when running on a single core and on four cores. Notably, the getCand function takes an order of magnitude more time compared to all the other functions. Furthermore, for mehenDi, we observed an increase in runtime as we raised the FDR threshold. The increase may be attributed to the exploration of more branches, as the threshold becomes less stringent. The parallelized version exhibits a similar pattern. We also observe the benefits of parallelization, achieving a more than threefold increase in speed using four cores compared to using a single core.

**Table 2:** Time taken in (hh:mm:ss) by treeclimbR and mehenDi on the BrSimNorm and MouseMuscle dataset.

| Method |  | BrSimNorm (hh:mm:ss) | MouseMuscle (hh:mm:ss) |
| --- | --- | --- | --- |
| `treeclimbR` | `getCand` | 37:06:05 | 15:05:40 |
| | `evalCand`($\alpha = 0.01$) | 00:00:51 | 00:00:28 |
| | `evalCand`($\alpha = 0.05$) | 00:00:50 | 00:00:29 |
| | `evalCand`($\alpha = 0.1$) | 00:00:50 | 00:00:29 |
| `mehenDi` (cores = 1) | $\alpha = 0.01$ | 00:07:16 | 00:06:49 |
| | $\alpha = 0.05$ | 00:08:49 | 00:12:14 |
| | $\alpha = 0.1$ | 00:09:60 | 00:15:23 |
| `mehenDi` (cores = 4) | $\alpha = 0.01$ | 00:01:58 | 00:02:22 |
| | $\alpha = 0.05$ | 00:02:25 | 00:03:55 |
| | $\alpha = 0.1$ | 00:02:46 | 00:04:44 |

## 3 Discussion

The high level of uncertainty that is observed in the abundances of some transcripts during RNA-Seq quantification poses a challenge for downstream analyses, particularly for tasks such as differential testing. Various differential testing methods have been proposed that incorporate uncertainty into testing, and these methods have been applied either to transcripts or genes. This approach can occasionally result in overlooking the signal for the features that are difficult to quantify. Our method mehenDi, makes use of the tree structure computed by `TreeTerminus` [Singh et al., 2023] to produce the selected nodes that are significant between conditions. mehenDi provides a data-driven procedure for finding signal at a resolution between the gene and transcript level for RNA-Seq differential analysis, where a selected inner node captures the signal for a set of underlying transcripts that have high uncertainty. While many tree-based differential testing procedures have been proposed, our method, mehenDi is among the few tailored toward RNA-Seq data, which requires specific considerations. Our method requires p-values for all the nodes of the tree and thus is flexible in the choice of the differential testing method used, though we have used Swish [Zhu et al., 2019] in this study.

We assessed and evaluated mehenDi on both the simulated and experimental datasets. In the variations of the simulated datasets without true signal, we observed that the FPR obtained by mehenDi was comparable to that generated by doing testing only on the transcripts. Similarly, on both variations of simulated data that included true signals, we observe an increased sensitivity for mehenDi while controlling the FDR at or very near the nominal level. The enhanced sensitivity is not only evident quantitatively through the metrics we report but is also reflected in the discovery of more transcript branches on the tree when conducting pairwise comparisons with other methods. Interestingly, `treeclimbR` [Huang et al., 2021], which has been designed to control FDR at the leaf level, shows a very high sensitivity but also outputs a large number of false positives. Similarly, on the experimental datasets, we observe many significant genes that do not contain a single underlying transcript that is deemed significant when doing transcript-level differential analysis, but which contribute to nodes reported by mehenDi. This is a reflection on the ability of mehenDi to find signals that would have been lost when doing transcript-level analysis. Shared exons are commonly observed among these transcripts that belong to the selected nodes, increasing our confidence in the utility of the approach. We also observed a clear reduction in uncertainty for the aggregated transcript groups compared to the individual transcripts. This can provide biologists the opportunity to narrow down to a set of transcripts for their analysis that belong to the transcript groups, which would have been lost when doing either transcript or gene-level analysis.

There exist several areas for improvement of our method, and also opportunities for developing new and related methods. The difference in the p-value distribution observed for the inner and leaf nodes in the null simulation when the p-values for all nodes are calculated together suggests the need for a differential testing method that takes the hierarchy of features into account. It seems that, at least in the different simulations that we have created, we are controlling the FDR. However, since we are using the same dataset to create the tree and do differential testing, this technically leads to double-dipping [Kriegeskorte et al., 2009], which can lead to an exaggeration of false positives. This could be one of the reasons why we observe an inflated FDR for `treeclimbR`.

To mitigate double-dipping, a Poisson/negative binomial sampling of the original data into independent partitions of the same dimensions was introduced in an approach called count splitting[Neufeld et al., 2022, 2023]. The first partition can be used for latent-variable estimation and the second for inference. Incorporating count splitting into mehenDi will be a promising future direction. The challenge, however, will be how to do this without negatively affecting the power, since mehenDi already appears empirically well-calibrated.

There might also be another source of false positives. When doing a multiple-hypothesis correction on a set of hypotheses, the error rate is controlled over the entire set and not on a subset. Using our filtering strategy, we are outputting a subset of hypotheses that have p-value below a certain threshold. Focused BH [Katsevich et al., 2023] provides a way of selecting a p-value threshold by running the filter for the different p-value thresholds and finding the largest threshold at which the computed conservative estimate of the false discovery proportion for the set is less than the desired error rate. The limitation of this approach is that, apart from being conservative, it also puts constraints on the properties of the filters. Further, Focused BH would be computationally slow in its native form, specifically if filters have to be computed over hundreds or thousands of p-value thresholds and each filter takes non-trivial time to run. Adapting this approach for mehenDi can also be explored in the future.

Finally, it would be intriguing to explore the development of a `TreeTerminus` and mehenDi like approach in the context of single-cell RNA-seq analysis, where there are quite distinct, but related, opportunities for data-driven aggregation to increase the power and specificity of differential analysis. While differential expression analysis has been the focus of this paper, an important future direction would also be to explore the usage of `TreeTerminus` trees and mehenDi for differential transcript usage (DTU) analysis.

## 4 Methods

### 4.1 Preliminaries

#### TreeTerminus

For a set of samples in an RNA-Seq experiment, `TreeTerminus` [Singh et al., 2023] outputs a forest of trees that summarizes the abundance uncertainty structure across them. The leaves of the individual trees comprise the set of quantified transcripts where each internal node represents an aggregation of the set of transcripts belonging to the subtree rooted on it, with no two trees having an overlapping set of transcripts/leaves. The mean uncertainty decreases as we ascend the trees. To enable downstream analysis, the trees in the forest are combined to obtain a single unified tree, T. The uncertainty of a node *n* for a sample *m* (leaf or inner) is estimated by the metric inferential relative variance (`infRV`)[Zhu et al., 2019] as:

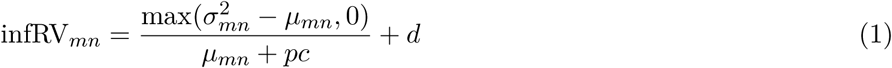

where *σ*^2^_mn_*, µ_mn_* are the variance and mean computed over the inferential replicates, *pc* is a pseudocount (with a default value of 5) and *d* is a small global shift (with a default value of 0.01). The inferential replicates refer to the bootstrap or Gibbs samples generated by `Salmon`. The meanInfRV for a node *n* is defined as the mean of infRV computed across the samples *m*. We describe `TreeTerminus` in more detail in Supplementary Section S1.1.

### 4.2 Problem Statement

For the given tree T, constructed on the set of *P* transcripts with all nodes including the leaves labelled as N = {1*, …, N* } and *N* ≥ *P*, we want to develop a method that outputs a set of **selected** nodes C ⊂ N which are differentially expressed between the conditions of interest. We want to leverage the tree to select nodes that may consist of both transcripts and inner nodes, with the nodes determined in a data-dependent manner to maximize the signal from the data while controlling for uncertainty. At the same time, we do not want to over-aggregate, especially if the features with signal exist at a lower level in the tree, since a finer resolution is always preferable. Further, any two nodes in the selected set should not share an ancestor/descendant relationship, i.e., for any two nodes *n_i_, n_j_* in C, Λ(*n_i_*) ∩ Λ(*n_j_*) = ∅, where Λ(*n_i_*) represents the set of transcripts that are the descendants of the node *n_i_* in the tree T. This ensures that a set of reads is not counted twice in the selected feature set. Our method mehenDi is designed taking the above principles into account. The inputs to mehenDi are the tree T, and the p-values for differential testing between the conditions, mean inferential relative variance (meanInfRV), and the direction of sign-change between conditions for all the nodes *N* belonging to T.

It is important to note that when mehenDi reports an inner node as selected, we do not imply anything about the differential status of its underlying descendant transcripts. The inner node should be interpreted as a standalone feature, the same way a gene is when doing gene-level analysis, where we are not always trying to comment on the differential status of its transcripts. Not all transcripts of a gene might be differentially expressed. A selected inner node might suggest that the signal existed in at least one underlying transcript, but was masked due to uncertainty.

### 4.3 mehenDi description

We will now outline our method, mehenDi. We first compute the p-values, meanInfRV and direction of sign change for all the nodes in the tree. Swish, which employs inferential replicates, is used to compute the p-values for the tree nodes. The counts for an inner node are obtained by aggregating the counts of the underlying transcripts. We recommend computing the p-values separately for the inner nodes and leaves. The direction of sign change is found by first computing the log fold change for a node across the biological samples at each inferential replicate individually and noting the direction (positive or negative) for each inferential replicate. If a direction emerges in a certain proportion of the inferential replicates beyond a threshold, the node is marked as confident, with that direction determining its sign change. Otherwise, the nodes are labelled as non-confident and are not assigned a direction of sign change. This is controlled by the parameter `minP`, which is set to 0.70 by default. Tree nodes with p-values below a specified threshold (pthresh) are identified and labelled as significant. The set of selected nodes, denoted as C, will be a subset of the significant nodes. The p-value threshold that is used for determining the significance of a node is computed by applying a multiple-testing procedure such as Benjamini-Hochberg (BH) [Benjamini and Hochberg, 1995] on the leaf nodes that controls the FDR on the leaves at rate *α*. mehenDi implements a top-down procedure starting from the root node, that first determines the branches which contain at least one significant node. Proceeding with this approach, it traverses each selected branch and, upon encountering a significant inner node, checks for the following criteria:

1. It has at least one non-significant node among its children.
2. All the descendant nodes that are marked confident have the same direction of sign change. This also means that all or a subset of descendant nodes can be non-confident.
3. All the child nodes have a meanInfRV above a certain threshold (`mIrvThresh`).

If all three criteria are met, that node is selected, and traversal along this branch terminates. Otherwise, this branch is traversed (recursively), checking the above criteria on each newly encountered significant node until either the node is selected or a leaf is reached. If a leaf is significant, it is added to the list of selected nodes. A small, illustrative example demonstrating the above algorithm is provided in Figure 1B.

#### 4.3.1 Explanation of the different criteria

The main intuition behind mehenDi is to aggregate transcripts in a data-driven manner to recover the signal that could have been lost due to uncertainty. The first criterion ensures a finer resolution of analysis since, if all children of a significant node are also significant, then the signal already exists at a lower level and aggregation is unnecessary. Owing to uncertainty, the direction of the sign change of a node can be ambiguous, as it might not be consistent across the inferential replicates. We thus assign a particular direction of sign change (positive/up, negative/down) to the nodes between the conditions of interest, only if we are confident. The direction and confidence of sign change of the descendant nodes should also have an impact in determining if a node is selected by mehenDi upstream. The second criterion helps to find a node having a consistent direction of the sign change across its underlying descendant nodes if they can be determined with certainty. This aids in having a clearer and more consistent biological interpretation of the node. The magnitude of effect size can also become smaller at a node if the underlying transcripts have an opposite direction of sign change and we should descend the tree to select nodes. The third criterion also checks for over-aggregation, since if we are certain about the abundance estimates for the nodes, we can confidently assess their differential signal and these nodes should not be aggregated. The default value of the parameter `mIrvThresh` is set to 0.40.

#### 4.3.2 Computing the p-values

`mehenDi` can take p-values produced by any differential analysis tool as input. However, in this study, we have used Swish [Zhu et al., 2019] for computing the p-values, since it takes inferential uncertainty into account. The p-values for the leaves and inner nodes are computed separately. The reasoning behind this choice is that the leaves have higher uncertainty compared to the inner nodes, especially the ones that appear higher in the tree (the height of a node refers to the maximum distance between the node and its underlying leaves). This leads to a relatively lower shrinkage of the test-statistic towards 0 for the inner nodes and increases the width of the distribution of the test-statistic used by Swish for computing the p-values for the inner nodes compared to those of the leaves. As a result, the p-values of the inner nodes will be smaller, on average, compared to the leaves, when the p-values are computed together.

### 4.4 Datasets

We evaluate mehenDi on four different datasets, two simulated and two experimental with differential signals, spanning different reference organisms with differential signals. In addition, we also evaluated mehenDi on simulated human datasets with no differential signal.

#### 4.4.1 Simulated human datasets

We used the pipeline defined in [Love et al., 2018] to create the simulated datasets. Polyester [Frazee et al., 2015] was used to generate the FASTQ files corresponding to the paired-end reads. Each simulated dataset consists of 12 samples belonging to 2 groups, with 6 samples in each group. TPM estimates were extracted from the GTEx V8 frontal cortex dataset to simulate samples with realistic ground-truth transcript-level expression. The TPM estimates were then used to create the count matrix (sim.counts.mat), which was provided as an input to Polyester to simulate the reads. The reads were simulated with a realistic fragment GC bias. The dispersion of transcript-level counts is drawn from the joint distribution of mean and dispersion values estimated from GEUVADIS samples [Lappalainen et al., 2013]. 80% of genes were set to be null genes, with no change in abundance between the two groups. All the transcripts for the next 10% of the genes were set to be differentially expressed, with all the transcripts belonging to a gene having the same fold change. For the remaining 10% of the genes, a single expressed transcript was selected to be differentially expressed. We created two variations of this simulation by varying the range of fold change: in the first variation, we keep the same range as in [Love et al., 2018] from 2 to 6; and in the second variation, the range is kept between 1.4 and 2.8. The first variation is referred to as `BrSimNorm` and the second as `BrSimLow`. The second simulation is created to measure the change in performance across the methods when the magnitude of effect sizes between the transcripts is decreased.

For the null simulations, we have a very similar setup with the key difference that the true fold change of all transcripts across conditions is set to 1.

#### 4.4.2 Mouse muscle dataset

We also analyzed a dataset taken from the skeletal muscle study GSE100505 [Terry et al., 2018]. We downloaded RNA-seq data for the 6 samples belonging to Masseter (MAST) and the 6 to Extensor digitorum longus (EDL) muscle tissues, with the accession numbers provided in Table S4. All the samples belong to the organism *Mus musculus*. This dataset is referred to as `MouseMuscle`.

#### 4.4.3 Chimpanzee brain dataset

The final dataset that has been analyzed in this study is the RNA-Seq data from [Sousa et al., 2017], with SynapseID syn7067053, collected from 5 Chimpanzees (*Pan Troglodyte*). We refer to this dataset as ChimpBrain. We specifically analyze samples obtained from the cerebellum and medial dorsal nucleus of thalamus brain tissues. The batch effects were observed for the specimen ID when visualizing the first two dimensions of the PCA obtained on the normalized counts matrix. We used svaseq [Leek, 2014] on the normalized counts using two surrogate variables to create a fit. The fit was then used to correct batch effects using the removeBatch-Effect function in limma [Ritchie et al., 2015], for each scaled and log-transformed inferential replicate. The exponential function was then applied to the batch-effect-corrected inferential replicates, converting counts back to the original scale.

### 4.5 Experimental Setup

#### 4.5.1 Pipeline to generate trees

For the ChimpBrain dataset, only BAM files were available as raw data, which were converted into FASTQ using the bamToFastq command from bedtools [Quinlan and Hall, 2010]. The quality control analysis for ChimpBrain and `MouseMuscle` datasets was done using fastqc [Andrews et al., 2012] and multiqc [Ewels et al., 2016]. `Salmon` indexes were built on gencode versions v26, vM25, and Pan tro 3.0 to analyze human, mouse and chimpanzee datasets [Frankish et al., 2019, Cunningham et al., 2022] respectively. `Salmon` was used to quantify and generate Gibbs replicates for the RNA-Seq samples. For each sample, 100 Gibbs replicates were generated using a thinning factor of 100. The trees from `TreeTerminus` were created using the Consensus mode. All the pipelines used for analysis in this paper were constructed using Snakemake [Mölder et al., 2021].

#### 4.5.2 Differential expression analysis

We compare the performance of carrying out the differential expression analysis using the different feature sets and tree-based methods on the analyzed datasets. The compared features comprise the selected transcripts (Txps), genes (Genes), Terminus groups (Terminus), and the best candidate, driver and selected nodes obtained by running the different tree-based methods on the unified tree T from `TreeTerminus`. The base set for Terminus comprises the discrete transcript groups and any remaining transcripts that belong to the transcript set of interest but are not covered by Terminus groups. For the tree-based methods, we compare mehenDi, `treeclimbR`, and BOUTH. Median-ratio scaling [Anders, 2010] was used to normalize the counts. When scaling the counts for the nodes in the tree, we first compute the size factor only using the leaf nodes and then divide the bias-corrected scaled counts of the inner nodes by the size factor. This procedure is described in detail in Supplementary Section S1.2. Swish is then used for generating p-values for the normalized counts. While `treeclimbR` and mehenDi require p-values for all the nodes in the tree, BOUTH requires the p-values only for the leaf nodes. The p-values were generated separately for the leaf and inner nodes using Swish. For `treeclimbR`, the parameter *α* is set to the nominal threshold, which aims to control the FDR at leaf nodes to extract the best candidates. We consider two feature sets for `treeclimbR` in our evaluation. `treeclimbR`(N) refers to the best candidate set obtained after running `treeclimbR`, and `treeclimbR`(L) consists of the descendant transcripts for the nodes in `treeclimbR`(N). We also consider `treeclimbR`(L) in our analysis because `treeclimbR` is designed to control FDR at the leaf level, which, in this case, corresponds to the transcripts.

For BOUTH, we set the parameter *FAR* to the nominal FDR, which controls for a false assignment rate, and we then extract the driver nodes from its output. A driver node provides the highest level of resolution with none of their ancestors being associated for differential analysis. Many of the descendant nodes of a driver node at different resolutions can be labelled as detected for differential analysis, which can complicate the analysis when trying to decide which resolution node to select, thus a driver node is chosen for our analysis. For mehenDi, the nominal FDR threshold is used to select the p-value threshold (on the leaves) to obtain the selected nodes. For the other features (Txps, Genes, Terminus), Swish is applied directly to their base sets individually, and the selected set consists of features that have their adjusted p-values less than the given nominal FDR threshold.

#### 4.5.3 Evaluation on the simulated datasets

Since the ground truth is known for the simulated datasets, we compute the True Positive Rate (TPR) and False Discovery Rate (FDR) on the selected features obtained from the different methods at the different nominal thresholds (0.01, 0.05, 0.1). When computing these metrics for the features that consist of transcript groups or the inner nodes in the tree, we only consider the status of the differential expression for that node or the transcript group in our evaluation. We do not evaluate the differential status at the underlying descendant transcripts or other inner nodes corresponding to the group or inner node. To obtain the ground truth for differential expression at an inner node, we first create the aggregated counts matrix for the tree T, sim.tree.counts.mat using sim.counts.mat. The sim.counts.mat consists of the true transcript counts corresponding to sample groups, on which we are evaluating the differential expression. This is the same matrix that was provided as an input to Polyester to simulate reads. The count of an inner node in sim.tree.counts.mat is computed by aggregating the counts of all the descendant transcripts in sim.counts.mat. The inner node is differentially expressed if the absolute log fold change (LFC) for that node computed using sim.tree.counts.mat is larger than a threshold. The threshold is set to the LFC obtained at the tree’s root node in the sim.tree.counts.mat.

## 5 Data Access

`mehenDi` is available as an R package and can be obtained from https://github.com/NPSDC/mehenDi. The scripts to reproduce the results in this manuscript are available at https://doi.org/10.5281/zenodo.11481255. The scripts to obtain and generate the datasets that have been analyzed in this manuscript are available at https://doi.org/10.5281/zenodo.11481348.

## 6 Contributions

N.P.S, M.I.L and R.P conceived the mehenDi method. N.P.S wrote the software. N.P.S, E.Y.W, M.I.L and R.P conceived the experiments, and N.P.S carried out the experiments. N.P.S, M.I.L and R.P interpreted the results. N.P.S, J.F carried out the simulations. N.P.S, M.I.L and R.P wrote and revised the manuscript. M.I.L and R.P secured funding.

