## Supplementary for "Tree-based differential testing using inferential uncertainty for RNA-Seq"

### S1 Methods

#### S1.1 TreeTerminus

For a given RNA-Seq experiment consisting of $M$ samples, `TreeTerminus` [4] outputs a forest of $K$ trees $\mathcal{T} = \{T_1, T_2, ..., T_K\}$, that summarize the abundance uncertainty structure across all the $M$ samples. The leaves of the individual trees comprise the set of quantified transcripts and each internal node represents an aggregation of the set of transcripts belonging to the subtree rooted at it, with no two trees having an overlapping set of transcripts/leaves. The input to `TreeTerminus` is the `salmon` [5] quantification estimates, $L$ inferential replicates, and range-factorized equivalence classes [3] corresponding to each RNA-Seq sample $m$, $m \in \{1, ..., M\}$. The $L$ inferential replicates are produced either through Gibbs sampling or bootstrap sampling and are denoted by $\mathcal{I}_{mi} = \{I_{mi_1}, I_{mi_2}, ..., I_{mi_L}\}$, where $I_{mi_l}$ represents the counts of the transcript $i$ for sample $m$ at the $l^{th}$ Gibbs/bootstrap iteration. The inferential replicate counts for an inner node are found by adding inferential replicate counts of each constituent transcript individually. The tree represents the order in which the different transcripts are aggregated into transcript groups starting from the leaf nodes and encoding different resolution layers for interpretation of the abundance of transcriptional groups, with uncertainty generally decreasing as one ascends the tree from the leaves. The uncertainty for any node (leaf or inner node) $n$ for a given sample $m$ is estimated using the metric `infRV` defined in [1], over $\mathcal{I}_{mn}$ as:

$$\text{infRV}_{mn} = \frac{\max(\sigma^2_{I_{mn}} - \mu_{I_{mn}}, 0)}{\mu_{I_{mn}} + pc} + d \tag{1}$$

where $\mu_{I_{mn}}, \sigma_{I_{mn}}$ are the mean, variance over the $L$ inferential replicates for a sample $m$ and node $n$, $pc$ is a pseudocount (with a default value of 5) and $d$ is a small global shift (with a default value of 0.01). The nodes situated at the lower heights in a branch in the tree usually represent the set of nodes for which large reduction in `infRV` was observed compared to its underlying children nodes. For most nodes in the tree, the underlying transcripts belong to the same gene, due to large sequence overlap between them which is a driving factor behind uncertainty. However, the transcripts in a node can also map to different genes, as there can be overlapping sequence regions between different genes as well, and also sequence similarity between genes belonging to a gene family. While a reduction in `infRV` governs how transcripts are aggregated into nodes, it does not mean that all the underlying transcripts will have similar strength and direction of differential signal between the conditions of interest.

**Unified Tree** - Once the forest from `TreeTerminus` is obtained, a unified tree is constructed. For the sake of simplicity, let $\mathcal{T}$ denote the unified tree. The unified tree is constructed using the `R` package `beaveR`(`https://github.com/COMBINE-lab/beaveR`). This tree is constructed by first creating a new root node and assigning all trees in the forest and the remaining transcripts in the transcriptome not covered by the trees as children of this root node.

### S1.2 Median-ratio scaled counts for the nodes in the tree

Using the formulation from [1], let $Y^0$ denote the counts matrix obtained for Salmon for the transcript set containing $M$ samples $1, .., m$ and $P$ transcripts $1, .., p$, with $Y^0_{ji}$ representing the counts for transcript $j$ in sample $i$. Let the matrix $Y^{\mathcal{T}0}$ denote the counts obtained for all the nodes in the tree $\mathcal{T}$ that has $P$ leaf nodes, where for an internal node $n$, $Y^{\mathcal{T}0}_{ni} = \sum_{d=1}^{|\Lambda(n)|} Y^0_{t_d i}, \forall t_d \in \Lambda(n)$, where $\Lambda(n)$ denote the indexes of the descendant transcripts of node $n$. The counts $Y^{\mathcal{T}}_{ji}$ are divided by a bias length term $b_{ji}$, accounting for the length w.r.t other transcripts:

$$b_{ji} = \frac{l_{ji}}{(\prod_{i=1}^{m} l_{ji})^{\frac{1}{m}}}$$

Then we divide the counts by $b_{ji}$ as

$$Y^{\mathcal{T}*}_{ji} = \frac{Y^{\mathcal{T}0}_{ji}}{b_{ji}}$$

The counts are then scaled to the geometric mean of sequencing depth as

$$Y_{ji}^{\mathcal{T}**} = \frac{Y_{ji}^{\mathcal{T}*}}{\sum_{j=1}^{P} Y_{ji}^{\mathcal{T}*}} \times \left( \prod_{i=1}^{m} \sum_{j=1}^{P} Y_{ji}^{\mathcal{T}0} \right)^{\frac{1}{m}}$$

For each sample $i$, a median-ratio size factor is computed as

$$s_i = \text{median}_j^P \frac{Y_{ji}^{\mathcal{T}**}}{\left( \prod_{k=1}^{m} Y_{jk}^{\mathcal{T}**} \right)^{\frac{1}{m}}}$$

We compute the size factor over only the leaf nodes. The final normalized counts are then computed as:

$$Y_{ji}^{\mathcal{T}} = \frac{Y_{ji}^{\mathcal{T}**}}{s_i}$$

The lengths of the inner nodes are computed using the strategy employed by `summarizeToGene` function in the `R` package `tximport` [2]. The length of an inner node $n$ for sample $i$ is computed as:

$$l_{ni}^{\mathcal{T}} = \frac{\sum\limits_{d=1}^{|\Lambda(n)|} l_{t_d i} tpm_{t_d i}}{\sum\limits_{d=1}^{|\Lambda(n)|} tpm_{t_d i}}, \forall t_d \in \Lambda(n)$$

Here *tpm* refers to transcripts per million estimates that are provided by Salmon.

### S1.3 Distance between nodes

The distance between the set of nodes $N_d$ and $N_p$ is computed as :

$$\begin{aligned}
\mathcal{D}(N_d, N_p) &= \frac{dist(N_d, N_p) + dist(N_p, N_d)}{2}, \\
dist(N_a, N_b) &= \frac{\sum\limits_{i=1}^{|N_a|} d(N_{ai})}{\|N_a\|}, \\
d(N_{ai}) &= \begin{cases}
0 & \text{if } N_{ai} \in N_b \\
Path_length(N_{ai}, N_{bk}) & \text{if } N_{bk} \in N_b \text{ and } N_{bk} \text{ is either an ancestor} \\
& \text{ or descendent of } N_{ai} \\
Path_length(N_{ai}, \text{root}) + 1 & \text{if } N_{ai} \notin N_b \text{ and no ancestor or} \\
& \text{descendant of } N_{ai} \text{ exists in } N_b
\end{cases}
\end{aligned}$$

where $N_d, N_p$ denote the node set output by `mehenDi` at the default parameters and parameter set $p$ respectively. $Path\_length(N_{ai}, N_{bk})$ denote the length of the path between the nodes $N_{ai}$ and $N_{bk}$. We are computing the average distance per node between the two sets. If the same node is present in both sets, the distance between them would be 0. Similarly, if for a node belonging to one set, there exists a node in the other set which is an ancestor/descendant for it, then the distance is computed by calculating the length of the path between them on the tree. On the other hand, if there is no ancestor/descendant for a node in the other set, then the distance is the length of the path from the root to that node with 1 added. 1 is added since this would be the lowest height node in the other set. The nodes that don't have an ancestor or descendant in the other set, can be the largest contributing factor to the distance metric and can create asymmetry for the overall distance metric aka $dist(N_a, N_b) \neq dist(N_b, N_a)$, as they do not directly have a counterpart in the other set. This can skew the metric, depending on the set w.r.t which distance is computed, especially if that set consists of nodes that represent unique branches in the tree. To balance this, our final distance metric $\mathcal{D}(N_d, N_p)$ is the average of $dist(N_d, N_p), dist(N_p, N_d)$.

[2] Charlotte Soneson, Michael I Love, Mark D Robinson. Differential analyses for RNA-seq: transcript-level estimates improve gene-level inferences. *F1000Research* 4, 2015.

[3] Mohsen Zakeri, Avi Srivastava, Fatemeh Almodaresi, Rob Patro. TreeTerminus—creating transcript trees using inferential replicate counts. *Bioinformatics* 33(14):i142–i151, 2016.

[4] Noor Pratap Singh, Michael I Love, Rob Patro. TreeTerminus—creating transcript trees using inferential replicate counts. *iScience* 26(6), 2023.

[5] Rob Patro, Geet Duggal, Michael I Love, Rafael A Irizarry, Carl Kingsford. Salmon provides fast and bias-aware quantification of transcript expression. *Nature methods* 14(4):417–419, 2017.

### S2 Tables

Table S1: Distribution of the height of the selected nodes output by `mehenDi` and candidate nodes by `treeclimbR` for the `BrSimNorm` dataset.

| Height of nodes | mehenDi | treeclimbR |
|---|---|---|
| 1 | 4013 | 2941 |
| 2 | 554 | 326 |
| 3 | 256 | 164 |
| 4 | 125 | 116 |
| 5 | 62 | 80 |
| 6-10 | 57 | 152 |
| >10 | 5 | 39 |
| Max height | 14 | 19 |

Table S2: Proportion of unique nodes that are observed for `mehenDi`, `treeclimbR` when doing a pairwise comparison for the `BrSimNorm` dataset at different nominal FDR. The number inside the parentheses represents the total number of nodes.

| Nominal FDR | 0.01 | 0.05 | 0.10 |
|---|---|---|---|
| `mehenDi` | 0.085 (199) | 0.412 (308) | 0.606 (381) |
| `treeclimbR(N)` | 0 (2) | 0 (4) | 0.25(4) |

Table S3: Proportion of unique nodes that are observed for `Txps`, `treeclimbR` when doing a pairwise comparison for the `BrSimNorm` dataset at different nominal FDR. The number inside the parentheses represents the total number of nodes.

| Nominal FDR | 0.01 | 0.05 | 0.10 |
|---|---|---|---|
| `Txps` | 0.103 (145) | 0.416 (255) | 0.590 (339) |
| `treeclimbR(N)` | 0 (15) | 0.24 (25) | 0.481 (27) |

Table S4: Accession IDs for the tissue samples for the `MouseMuscle` dataset that have been used for analysis in this paper.

| **Accession ID** | **TissueName** |
|---|---|
| SRR5758666 | Mast |
| SRR5758667 | Mast |
| SRR5758668 | Mast |
| SRR5758669 | Mast |
| SRR5758670 | Mast |
| SRR5758671 | Mast |
| SRR5758630 | EDL |
| SRR5758631 | EDL |
| SRR5758632 | EDL |
| SRR5758633 | EDL |
| SRR5758634 | EDL |
| SRR5758635 | EDL |

Table S5: Number of differentially expressed genes obtained at the different nominal thresholds for the `MouseMuscle` dataset between EDL and MAST.

| Nominal FDR | 0.01 | 0.05 | 0.10 |
|---|---|---|---|
| Number of DEGs | 2535 | 4494 | 5814 |

Table S6: Number of differentially expressed transcripts obtained at the different nominal FDR thresholds for the `MouseMuscle` dataset between EDL and MAST.

| Nominal FDR | 0.01 | 0.05 | 0.10 |
|---|---|---|---|
| Number of DTEs | 3149 | 6849 | 9592 |

Table S7: Distribution of the height of the `mehenDi` nodes for the `MouseMuscle` dataset.

| Nominal FDR | 0.01 | 0.05 | 0.10 |
|---|---|---|---|
| 1 | 2522 | 5406 | 7500 |
| 2 | 651 | 1285 | 1698 |
| 3 | 295 | 555 | 696 |
| 4 | 134 | 252 | 308 |
| $\geq 5$ | 109 | 195 | 256 |

Table S8: Distribution of the total number of unique genes, the transcripts covered by `mehenDi` nodes map to for the `MouseMuscle` dataset.

| Nominal FDR | 0.01 | 0.05 | 0.10 |
|---|---|---|---|
| 1 | 3672 | 7625 | 10369 |
| 2 | 33 | 56 | 73 |
| 3 | 3 | 8 | 9 |
| 4 | 0 | 4 | 4 |
| 5 | 1 | 0 | 0 |
| 7 | 0 | 0 | 1 |
| 9 | 0 | 0 | 2 |

Table S9: Number of differentially expressed genes (DEGs) that don't contain a single underlying transcript that is called significant at the different nominal FDR thresholds for the `MouseMuscle` dataset.

| Nominal FDR | 0.01 | 0.05 | 0.10 |
|---|---|---|---|
| Number of DEGs | 685 | 839 | 907 |

Table S10: Number of differentially expressed transcripts (DETs) that don't map to a single significant gene at the different nominal FDR thresholds for the `MouseMuscle` dataset.

| Nominal FDR | 0.01 | 0.05 | 0.10 |
|---|---|---|---|
| Number of DETs | 835 | 1736 | 2214 |

Table S11: Number of `mehenDi` nodes that neither map to a significant gene nor contain a single significant transcript for the `MouseMuscle` dataset.

| Nominal FDR | 0.01 | 0.05 | 0.10 |
|---|---|---|---|
| Number of mehenDi nodes | 231 | 382 | 390 |

Table S12: Number of differentially expressed genes (DEGs) at the different nominal FDR thresholds for the `ChimpBrain` dataset.

| Nominal FDR | 0.01 | 0.05 | 0.10 |
|---|---|---|---|
| Number of DEGs | 6090 | 8256 | 9354 |

Table S13: Number of differentially expressed transcripts (DETs) at the different nominal FDR thresholds for the `ChimpBrain` dataset.

| Nominal FDR | 0.01 | 0.05 | 0.10 |
|---|---|---|---|
| Number of DETs | 3539 | 10133 | 11886 |

Table S14: Distribution of the height of the `mehenDi` nodes for the `ChimpBrain` dataset.

| Nominal FDR | 0.01 | 0.05 | 0.10 |
|---|---|---|---|
| 1 | 2721 | 7979 | 9272 |
| 2 | 705 | 1387 | 1731 |
| 3 | 342 | 554 | 643 |
| 4 | 126 | 196 | 229 |
| $\geq 5$ | 107 | 131 | 142 |

Table S15: Distribution of the total number of unique genes, the transcripts covered by `mehenDi` nodes map to for the `ChimpBrain` dataset.

| Nominal FDR | 0.01 | 0.05 | 0.10 |
|---|---|---|---|
| 1 | 3943 | 58 | 11842 |
| 2 | 10106 | 140 | 1 |
| 3 | 11842 | 174 | 1 |

Table S16: Number of differentially expressed genes (DEGs) that don't contain a single underlying transcript that is called significant at the different nominal FDR thresholds for the `ChimpBrain` dataset.

| Nominal FDR | 0.01 | 0.05 | 0.10 |
|---|---|---|---|
| Number of DEGs | 2984 | 787 | 716 |

Table S17: Number of differentially expressed transcripts (DETs) that don't map to significant genes for the `ChimpBrain` dataset.

| Nominal FDR | 0.01 | 0.05 | 0.10 |
|---|---|---|---|
| Number of DETs | 130 | 835 | 854 |

Table S18: Number of `mehenDi` nodes that neither map to a significant gene nor contain a single significant transcript for the `ChimpBrain` dataset.

| Nominal FDR | 0.01 | 0.05 | 0.10 |
|---|---|---|---|
| Number of mehenDi nodes | 31 | 59 | 76 |

### S3 Figures

Figure S1: Distribution of the p-values for the leaf and inner nodes on the null simulations, when the hypothesis testing is carried out separately.

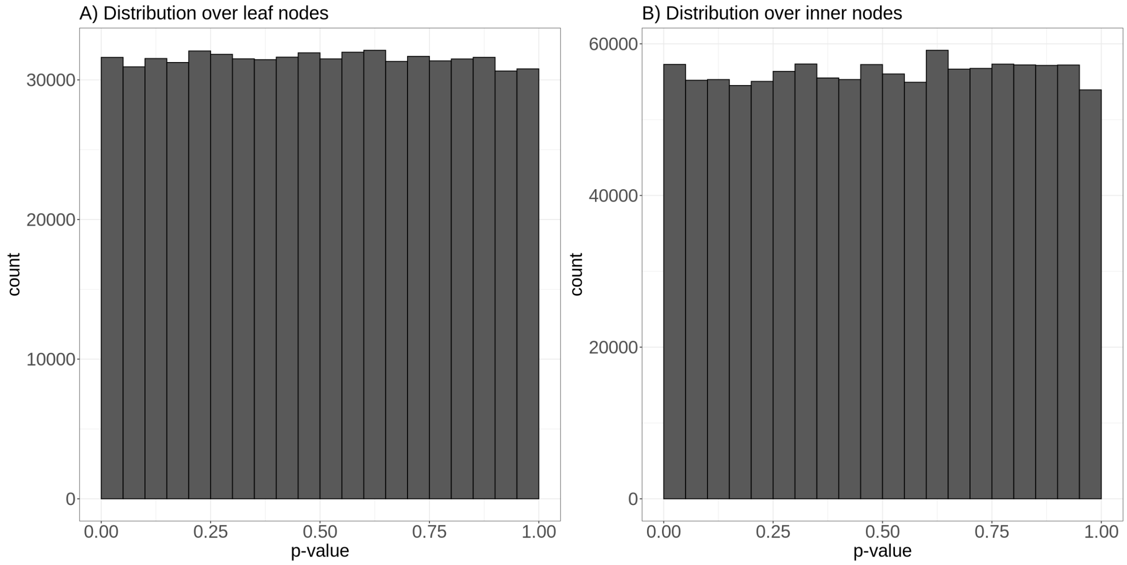


Figure S2: UpSet plot covering the number of true positive transcripts that are covered by the nodes that are output by the different methods for the `BrSimNorm` dataset nominal FDR.

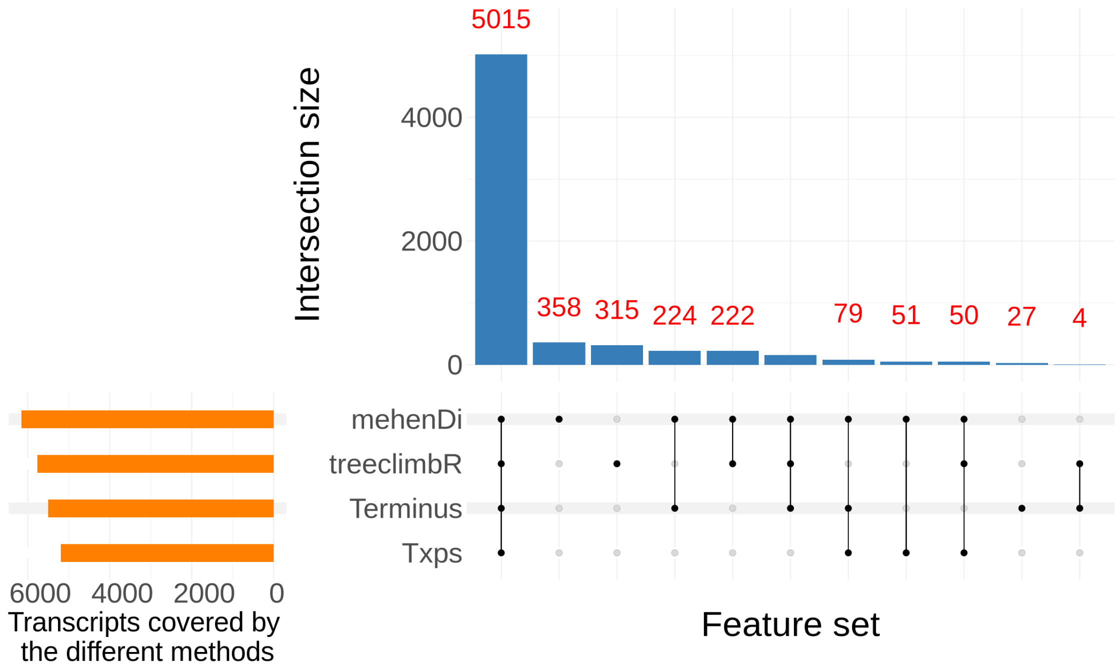

Figure S3: UpSet plot covering the number of true positive transcripts that are covered by the nodes that are output by the different methods for the `BrSimLow` dataset at the 0.01 nominal FDR.

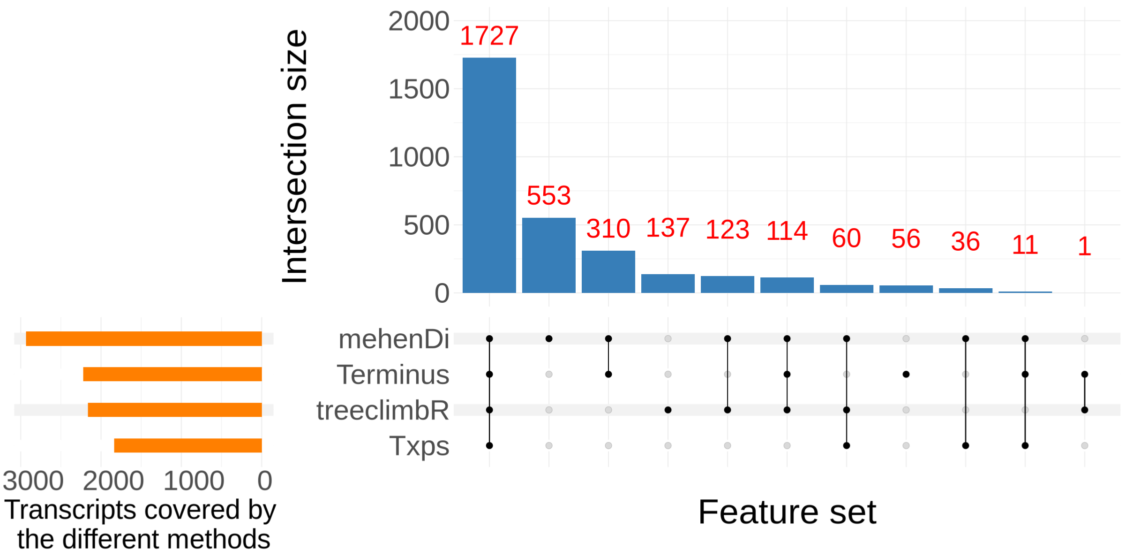


Figure S4: True Positive Rates and Empirical False Discovery Rates at the different nominal FDR thresholds by individually varying the parameters `minP` and `mIrvThresh` for the `BrSimNorm` dataset. Both the metrics have been rounded to 3 decimal places

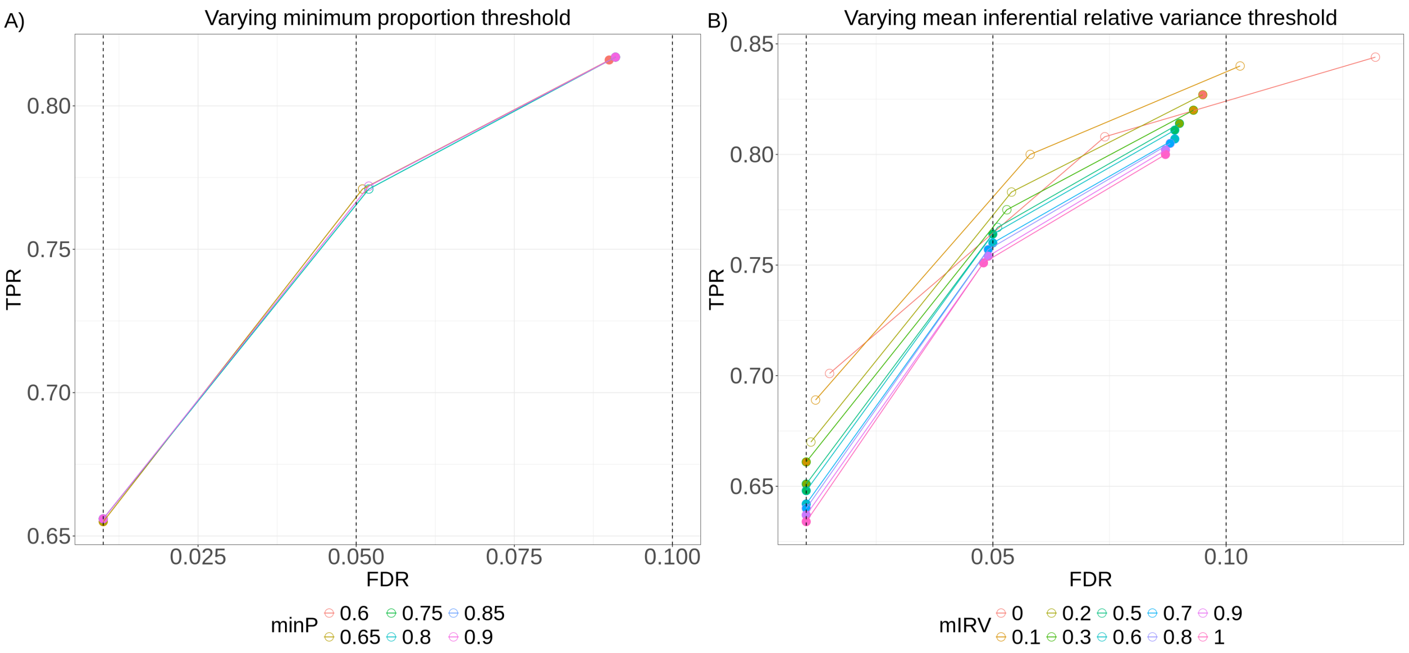

Figure S5: Example of a `mehenDi` node that is overaggregated in `Terminus` for the `BrSimNorm` dataset. A) Subtree representing the transcripts covered by the `Terminus` group. B) Inferential replicates for the `Terminus` group. C) Inferential replicates for the selected node output by `mehenDi`.

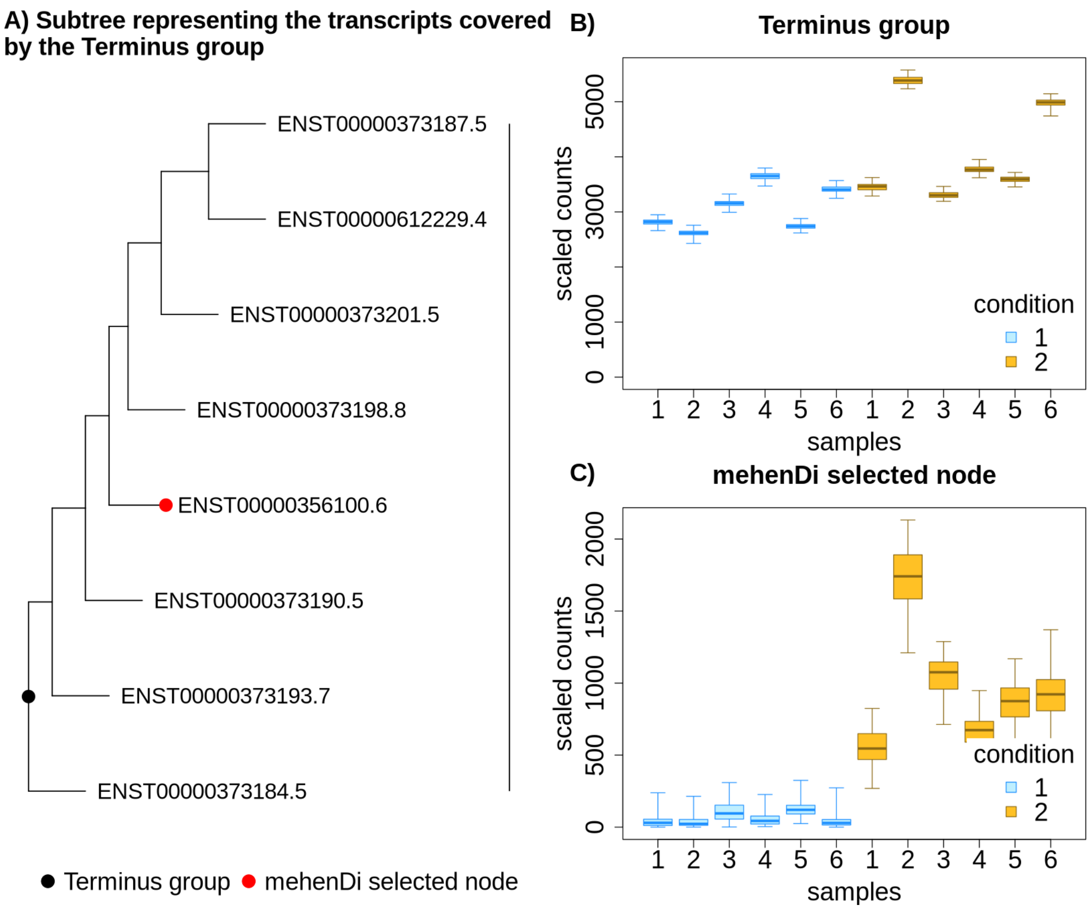

Figure S6: Example of a `mehenDi` node that is overaggregated in `Terminus` for the `BrSimNorm` dataset. A) Subtree representing the transcripts covered by the `Terminus` group. B) Inferential replicates for the `Terminus` group. C) Inferential replicates for the selected node output by `mehenDi`.

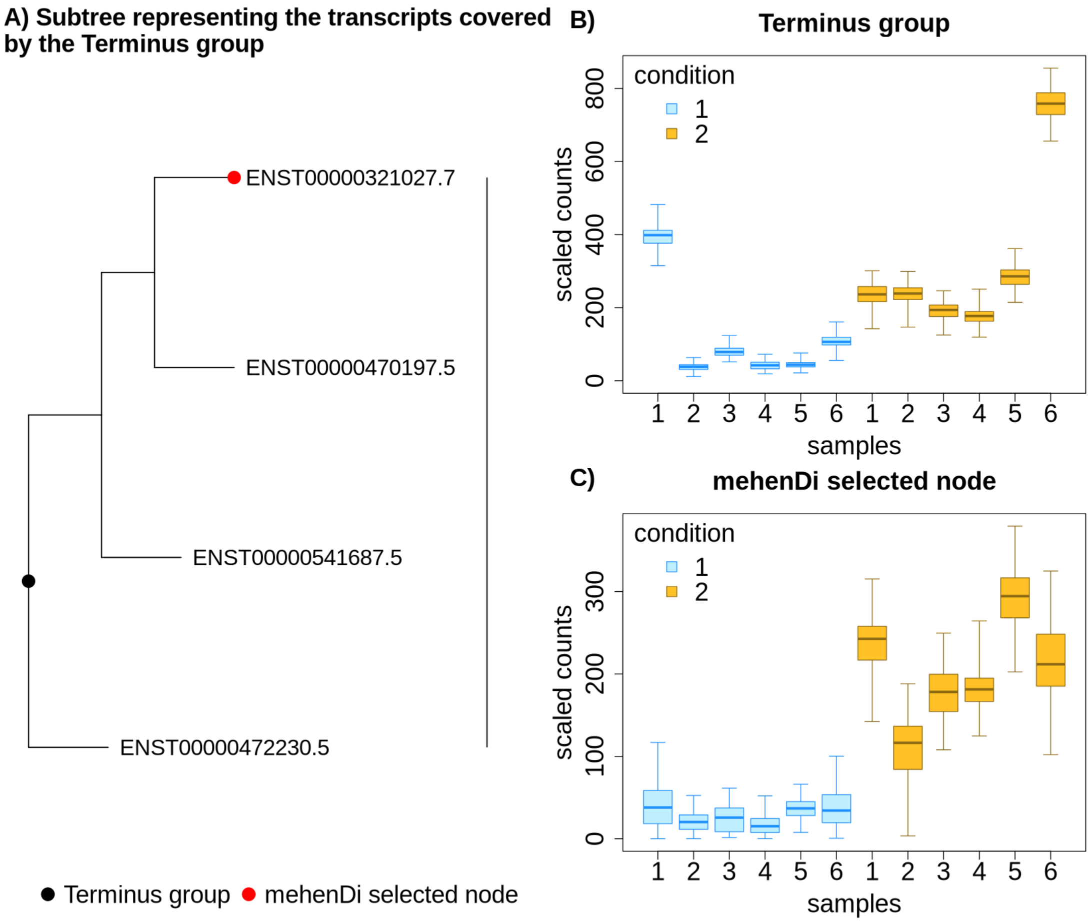

Figure S7: Example of a `mehenDi` node that is not aggregated enough in `Terminus` for the `BrSimNorm` dataset. A) Subtree representing the transcripts covered by the `mehenDi` group. B) Inferential replicates for the `Terminus` group. C) Inferential replicates for the selected node output by `mehenDi`.

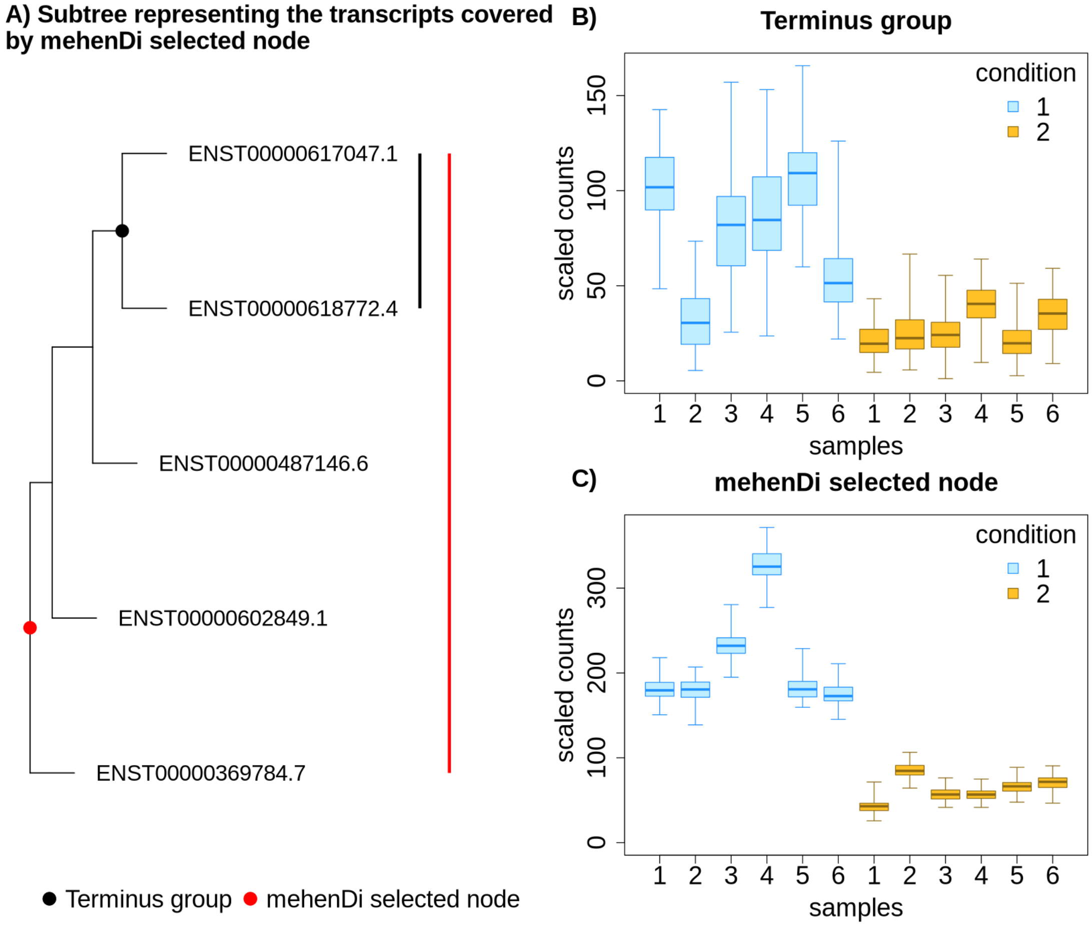

Figure S8: Example of a `mehenDi` node that is not aggregated enough in `Terminus` for the `BrSimNorm` dataset. A) Subtree representing the transcripts covered by the `mehenDi` group. B) Inferential replicates for the `Terminus` group. C) Inferential replicates for the selected node output by `mehenDi`.

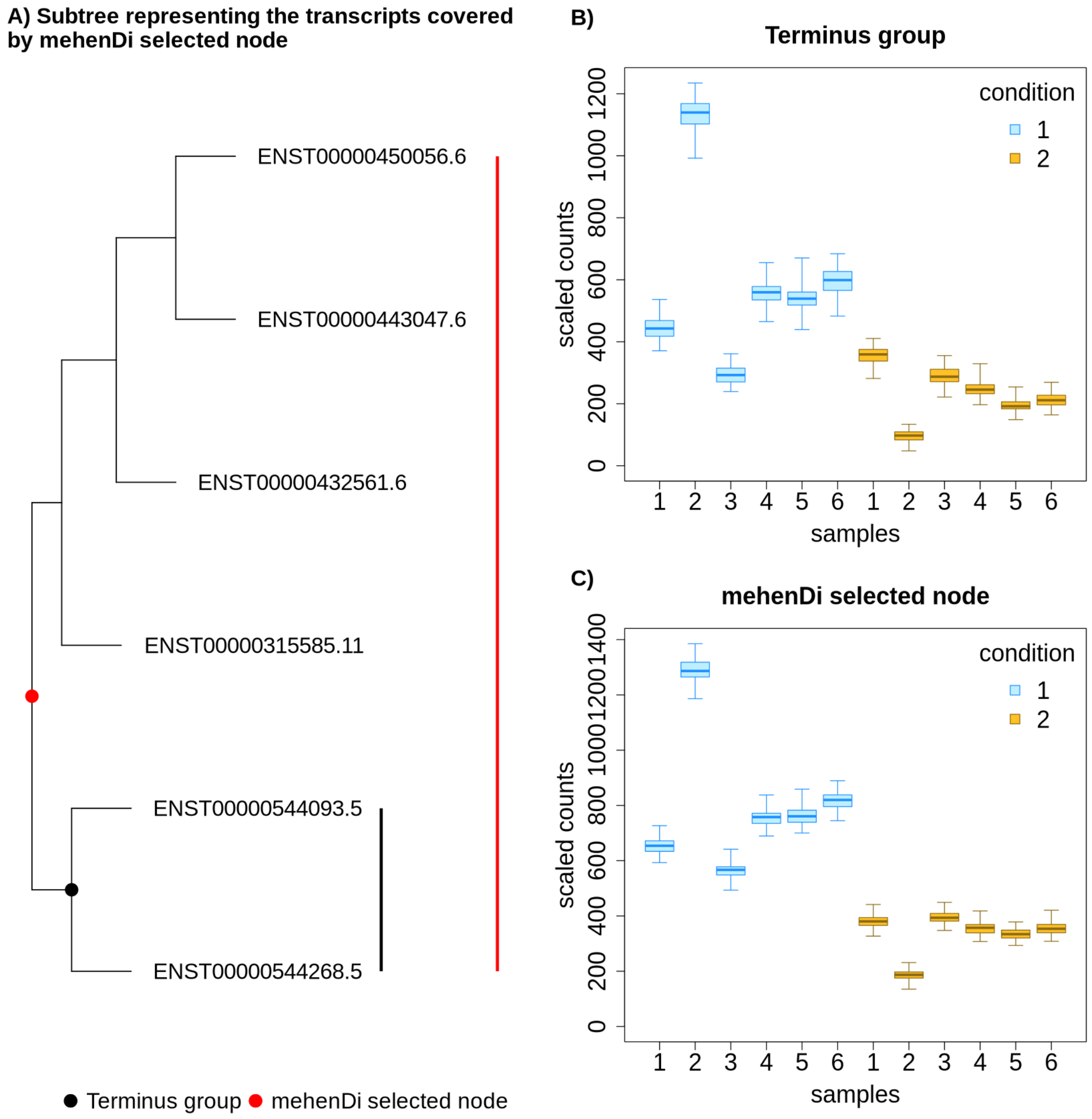

Figure S9: Examination of error metrics for the unique nodes obtained for `treeclimbR` when doing `treeclimbR` vs `Txps` analysis at the different nominal FDR thresholds. We vary the magnitude of log fold change (LFC) and plot the empirical FDR and the total number of nodes that are left after filtering the unique nodes based on LFC.

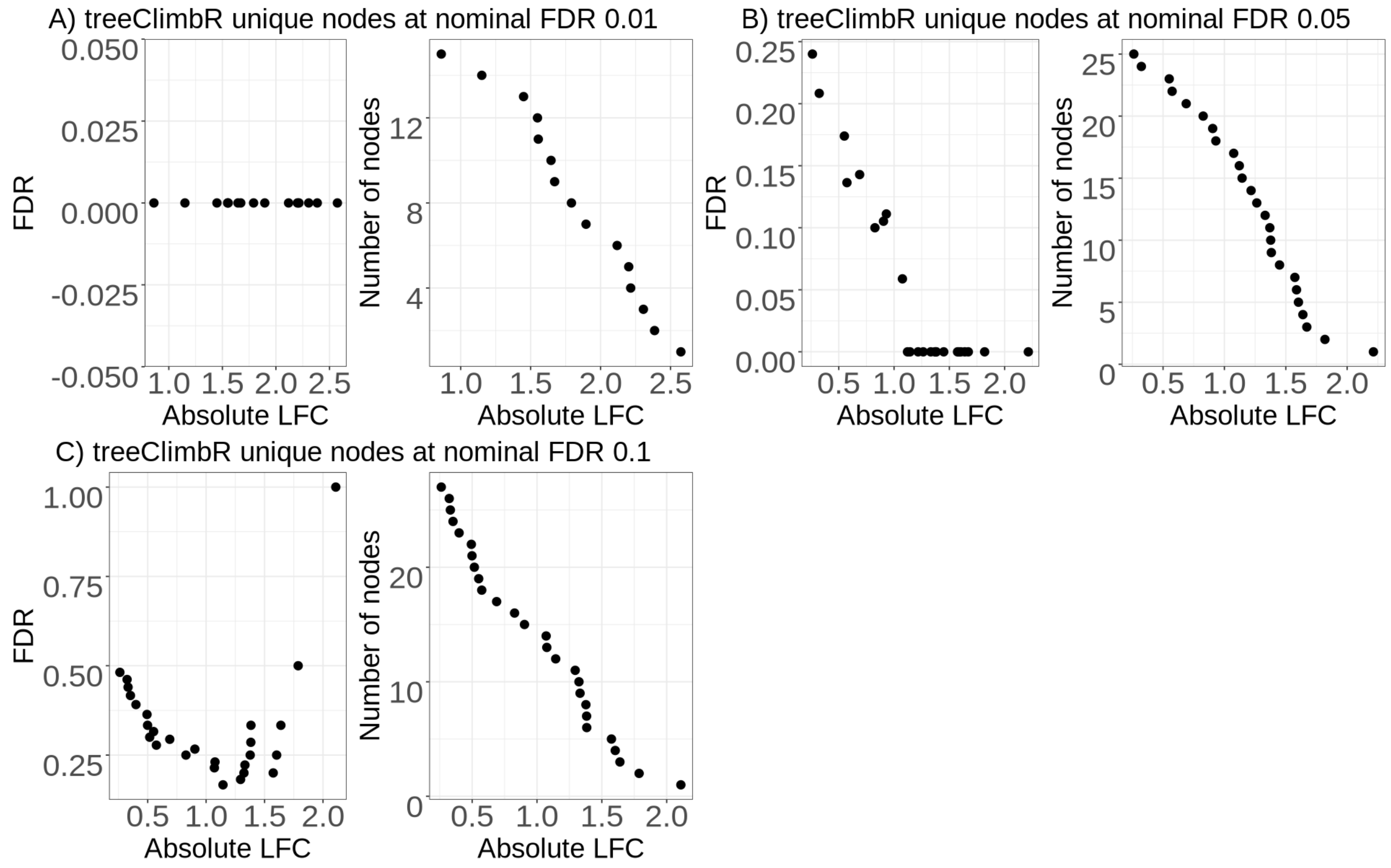

Figure S10: Examination of error metrics for the unique nodes obtained for `Txps` when doing `treeclimbR` vs `Txps` analysis at the different nominal FDR thresholds. We vary the magnitude of log fold change (LFC) and plot the empirical FDR and the total number of nodes that are left after filtering the unique nodes based on LFC.

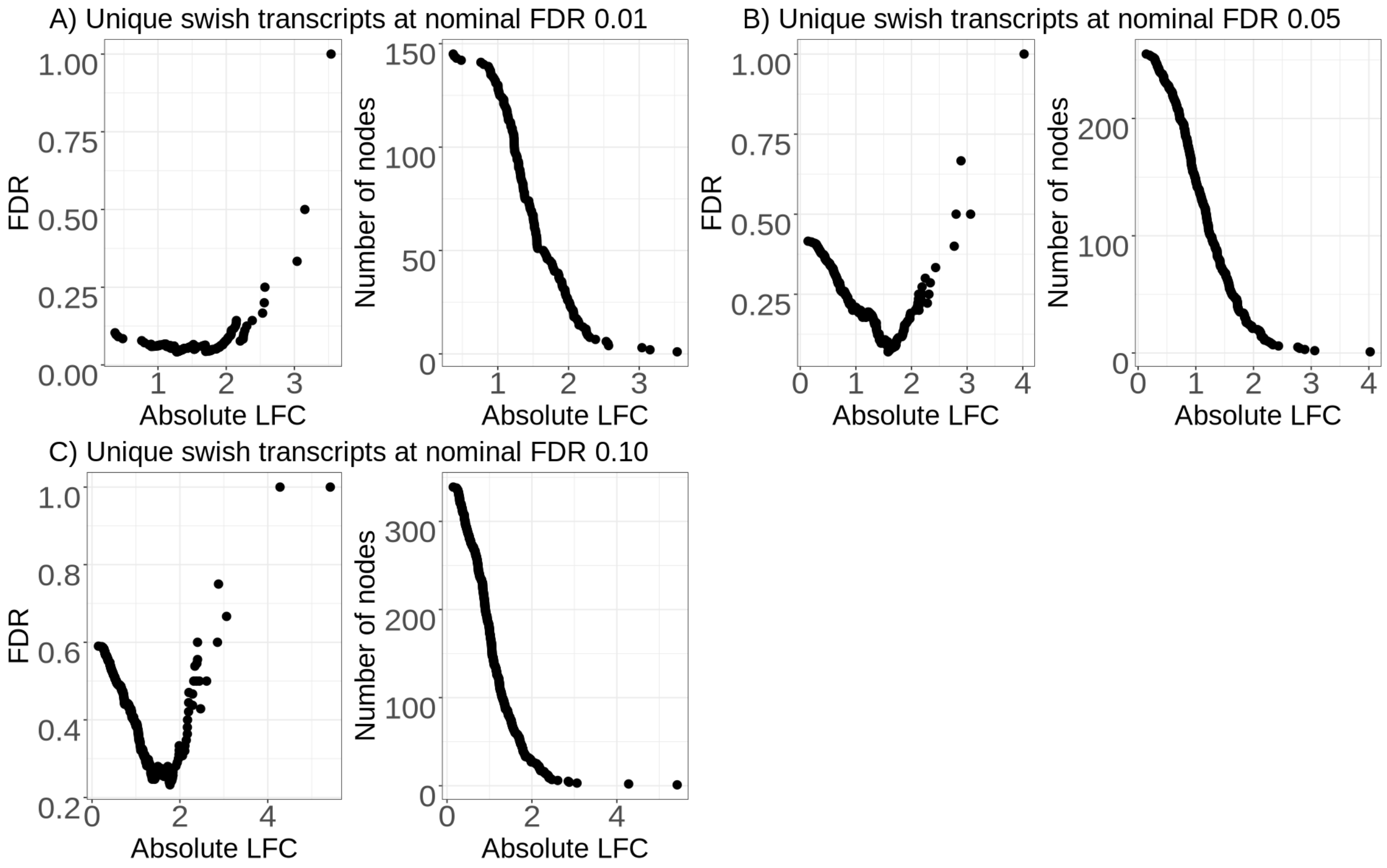

Figure S11: The first two dimensions of the PCA using the top 1000 variable features for the `MouseMuscle` dataset

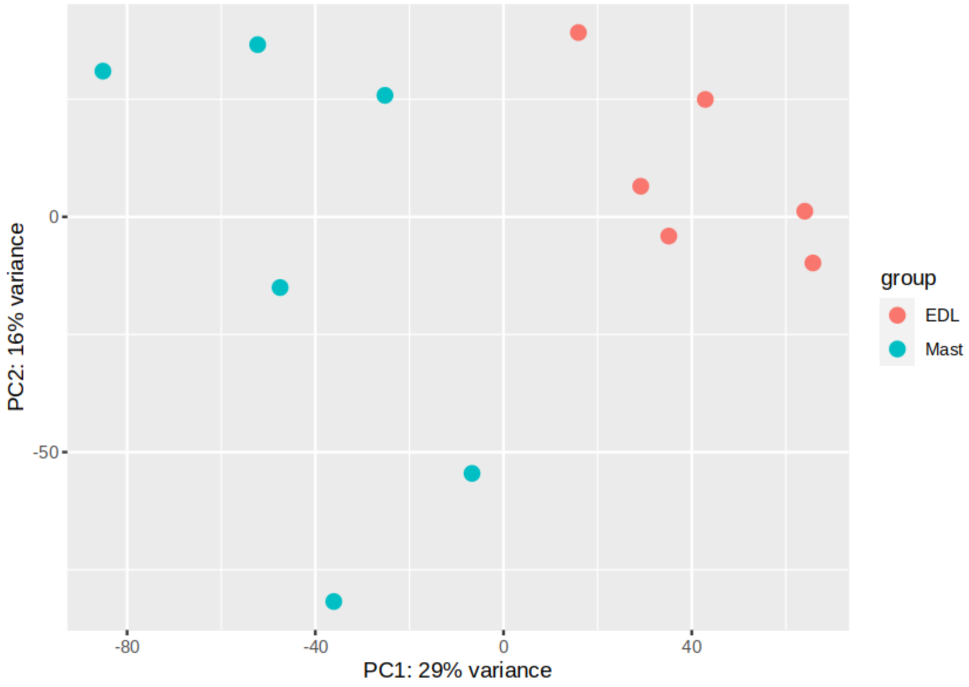

Figure S12: The average distance between the nodes obtained for `mehenDi` using default parameters and varying `minP` and `mIrvThresh` individually for the `MouseMuscle` dataset.

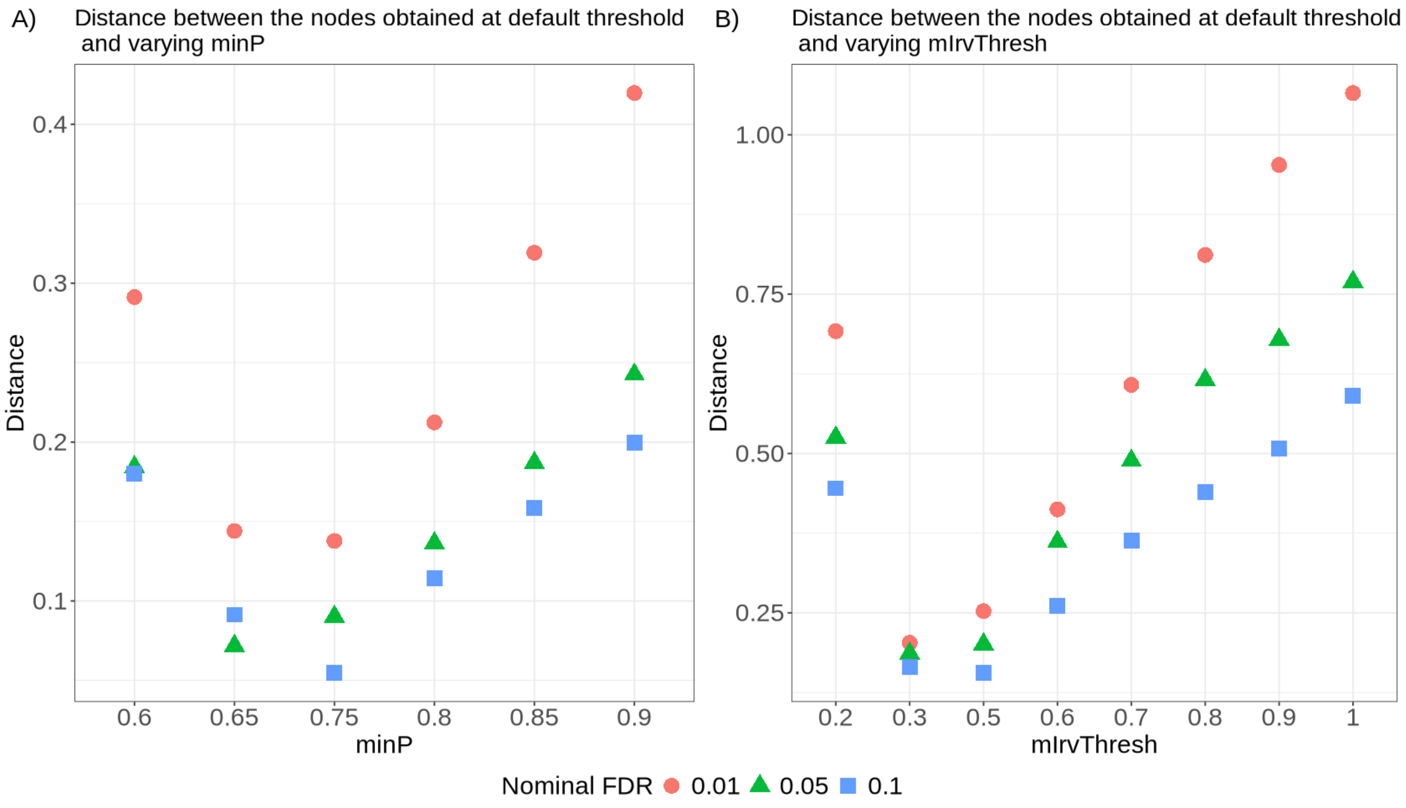


Figure S13: UpSet plot covering the number of transcripts that are covered by the nodes that are output by the different methods for the `MouseMuscle` dataset.

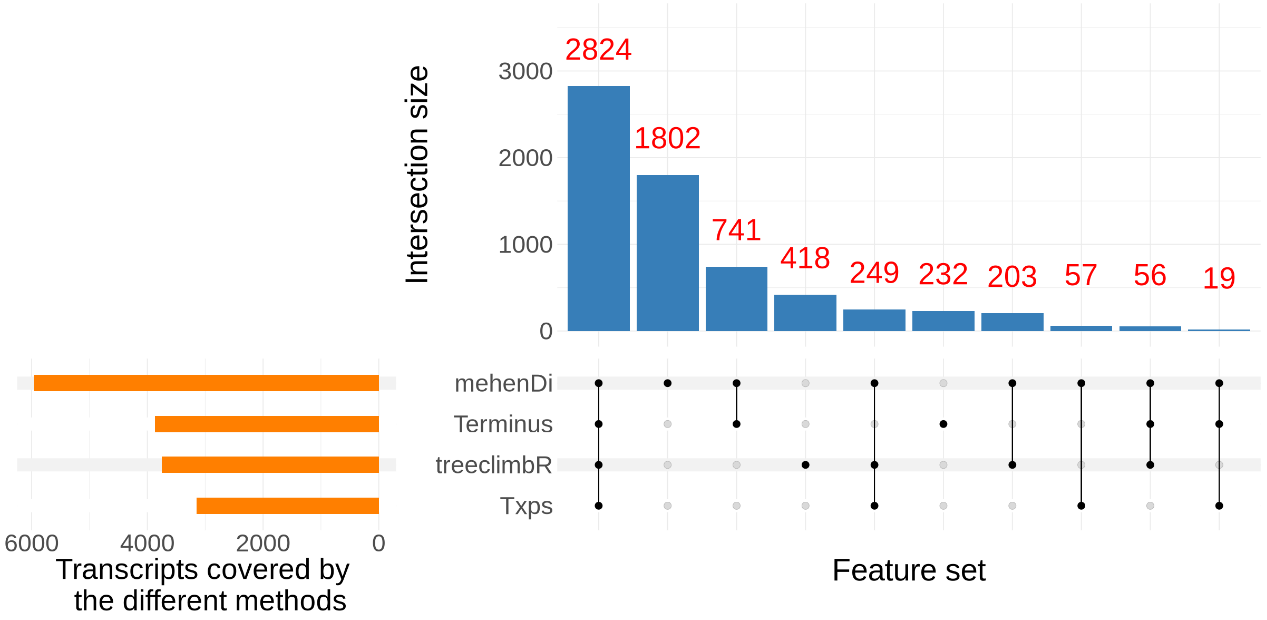

Figure S14: Examining the transcript profile for the gene *Hmcn2* in the `MouseMuscle` dataset. A) Transcripts in a pileup style. B) Tree representing the transcripts covered by the gene *Hmcn2*, with the red node representing the transcripts covered by the `mehenDi` selected node. C) Inferential replicates for the transcript ENSMUST00000138821.7, which had the lowest p-value among all the transcripts in the tree. D) Inferential replicates for the `mehenDi` selected node.

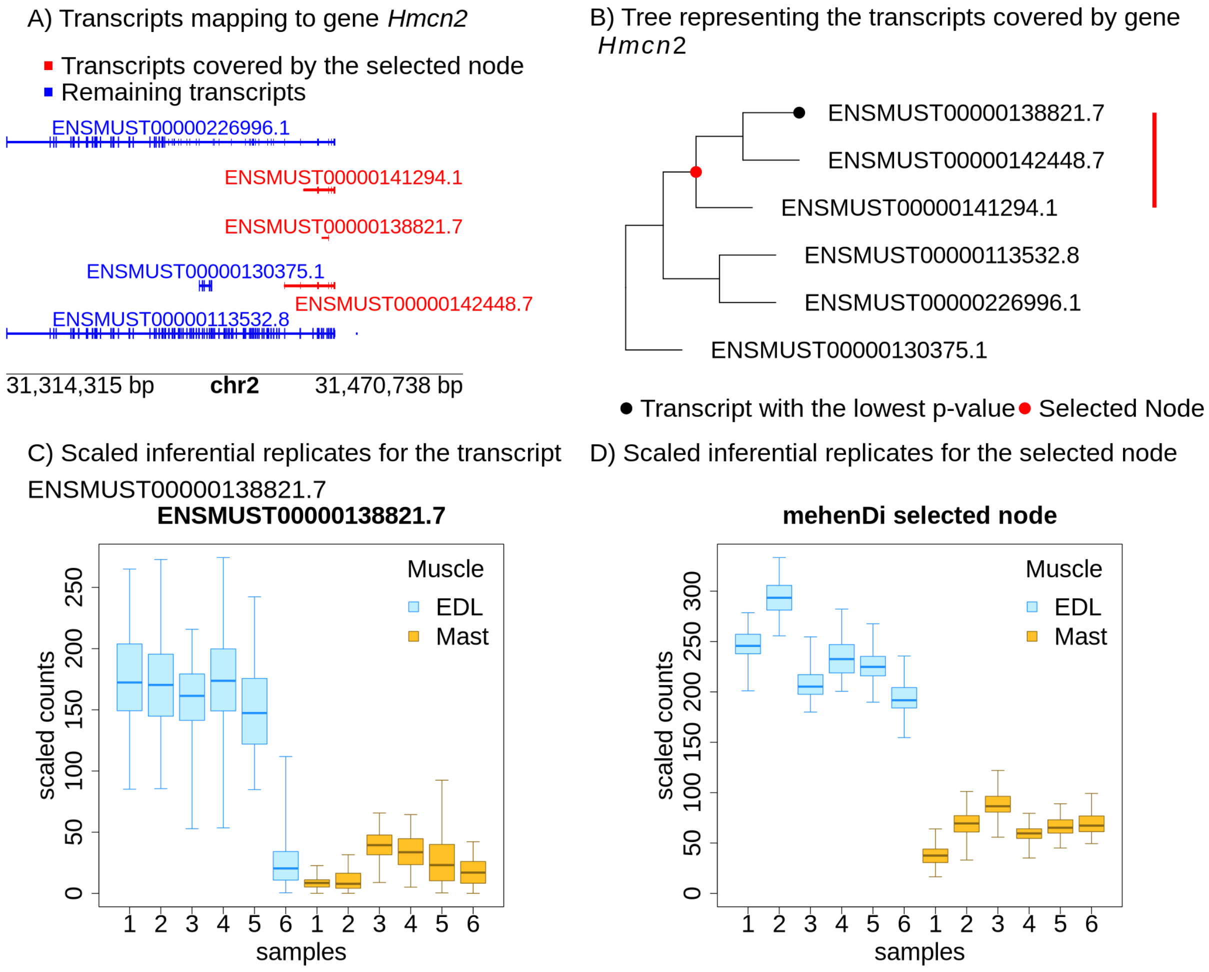

Figure S15: Examining the transcript profile for the gene *Emid1* in the `MouseMuscle` dataset. A) Transcripts in a pileup style. B) Tree representing the transcripts covered by the gene *Emid1*, with the red node representing the transcripts covered by the `mehenDi` selected node. C) Inferential replicates for the transcript ENSMUST00000151906.7, which had the lowest p-value among all the transcripts in the tree. D) Inferential replicates for the `mehenDi` selected node.

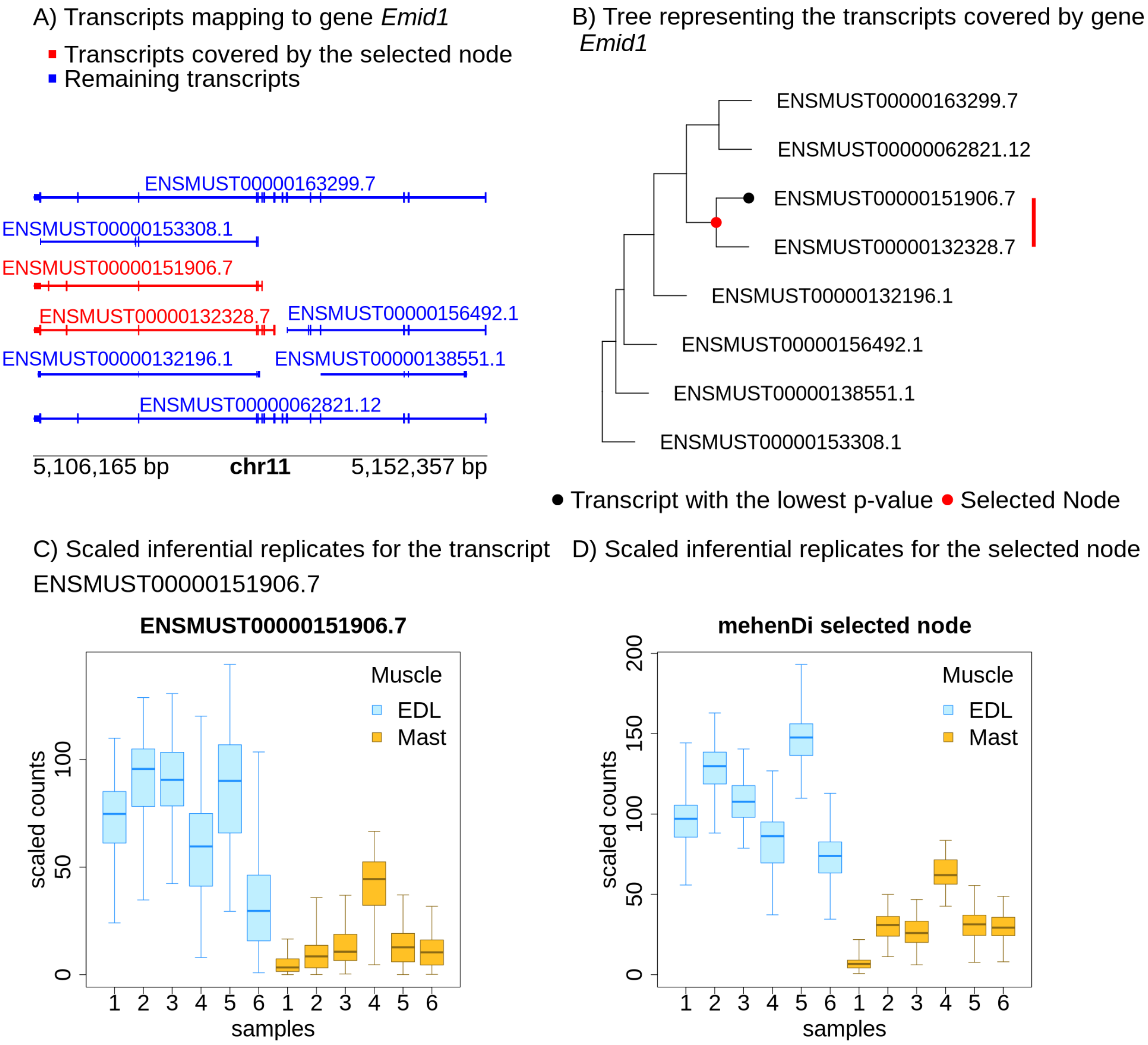

Figure S16: Examining the transcript profile for the gene *Prss55* in the `MouseMuscle` dataset. A) Transcripts in a pileup style. B) Tree representing the transcripts covered by the gene *Prss55*, with the red node representing the transcripts covered by the `mehenDi` selected node. C) Inferential replicates for the transcript ENSMUST00000171503.7, which had the lowest p-value among all the transcripts in the tree. D) Inferential replicates for the `mehenDi` selected node.

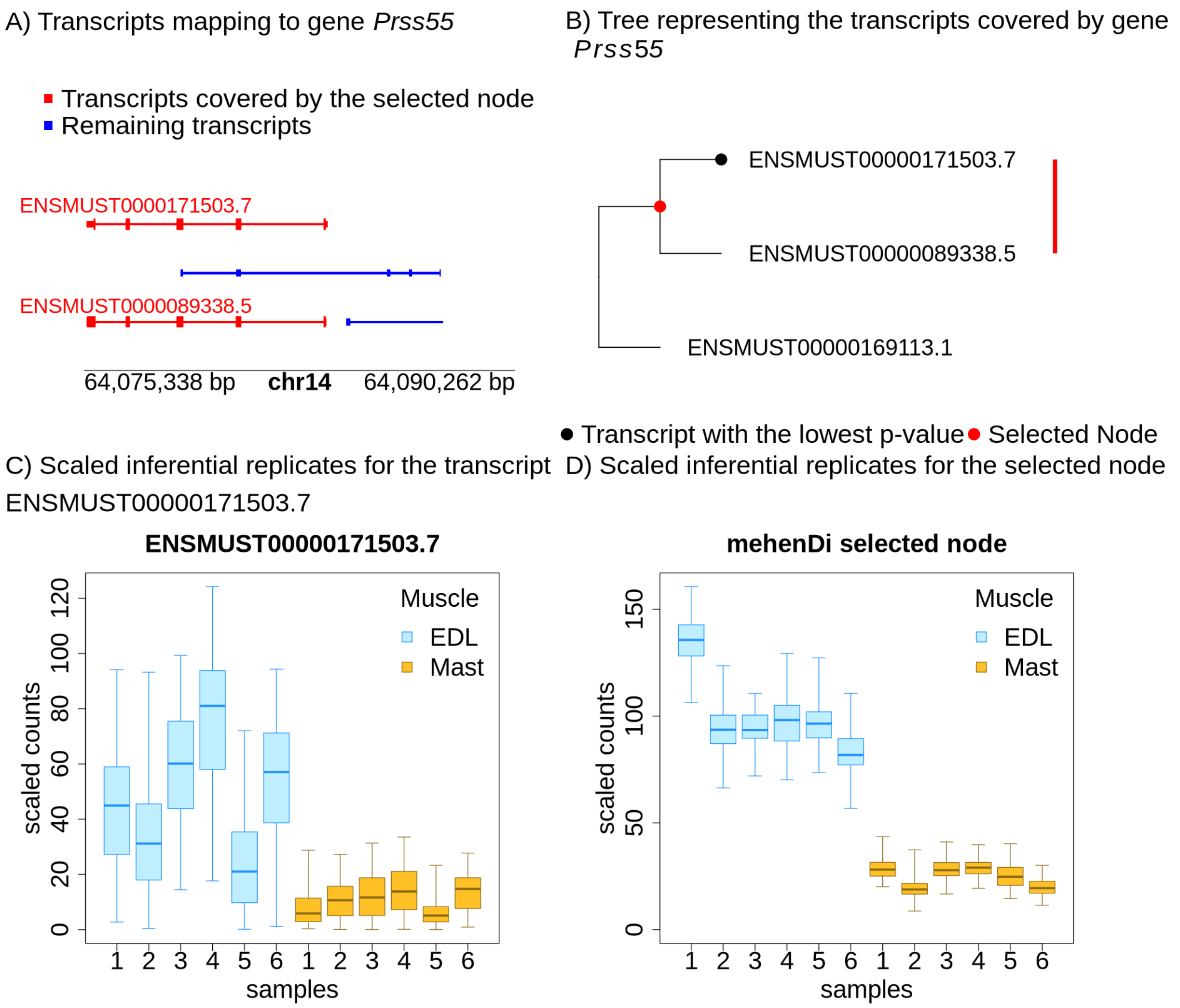

Figure S17: Examining the transcript profile for the gene *Stard10* in the `MouseMuscle` dataset. A) Transcripts in a pileup style. B) Tree representing the transcripts covered by the gene *Stard10*, with the red node representing the transcripts covered by the `mehenDi` selected node. C) Inferential replicates for the transcript ENSMUST000000032927.13, which had the lowest p-value among all the transcripts in the tree. D) Inferential replicates for the `mehenDi` selected node.

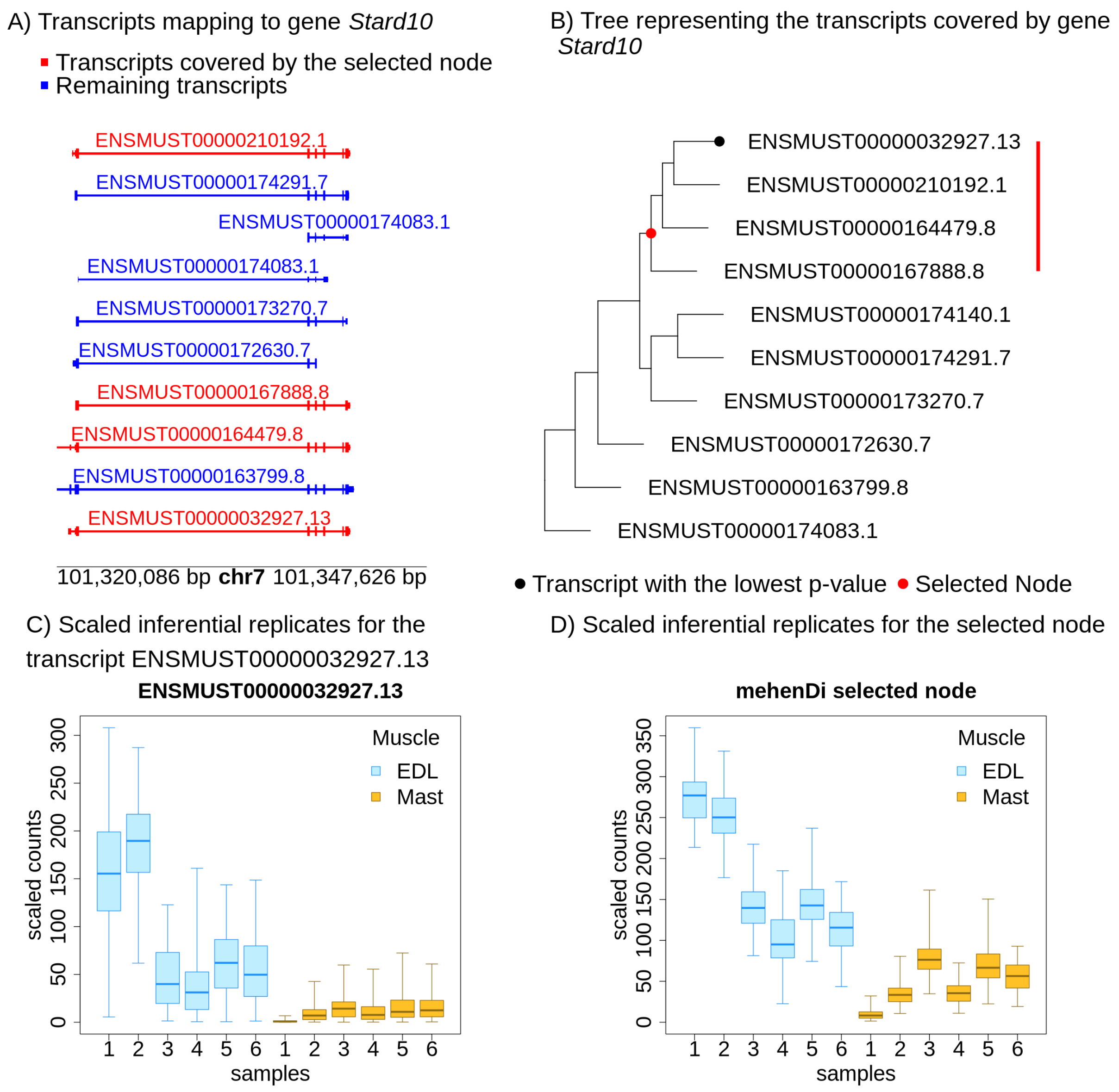

Figure S18: Examining the transcript profile for the gene *Syk* in the `MouseMuscle` dataset. A) Transcripts in a pileup style. B) Tree representing the transcripts covered by the gene $SYK$, with the red node representing the transcripts covered by the `mehenDi` selected node. C) Inferential replicates for the transcript ENSMUST00000055087.6, which had the lowest p-value among all the transcripts in the tree. D) Inferential replicates for the `mehenDi` selected node.

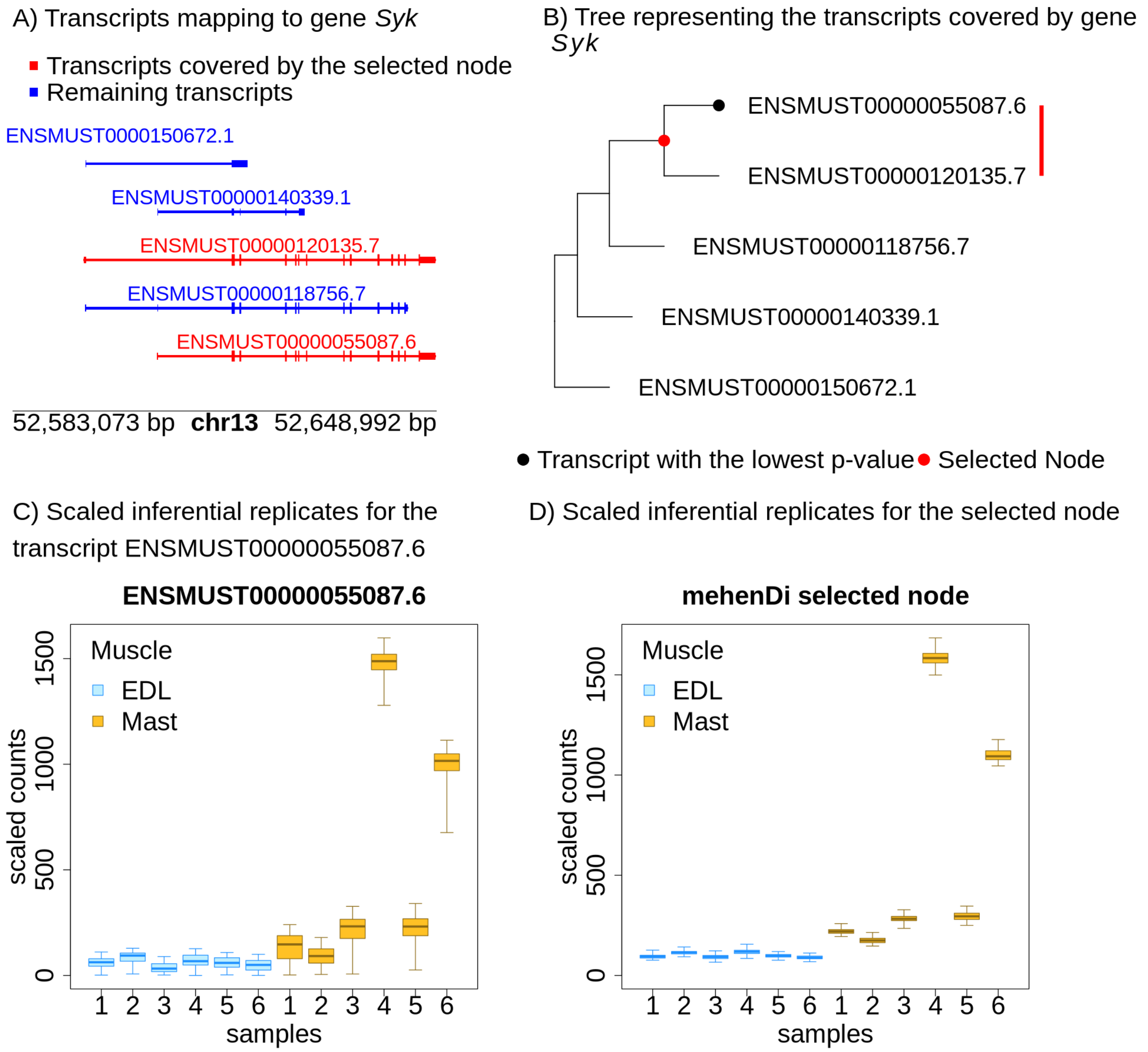

Figure S19: UpSet plot covering the number of transcripts that are covered by the nodes that are output by the different methods for the `ChimpBrain` dataset.

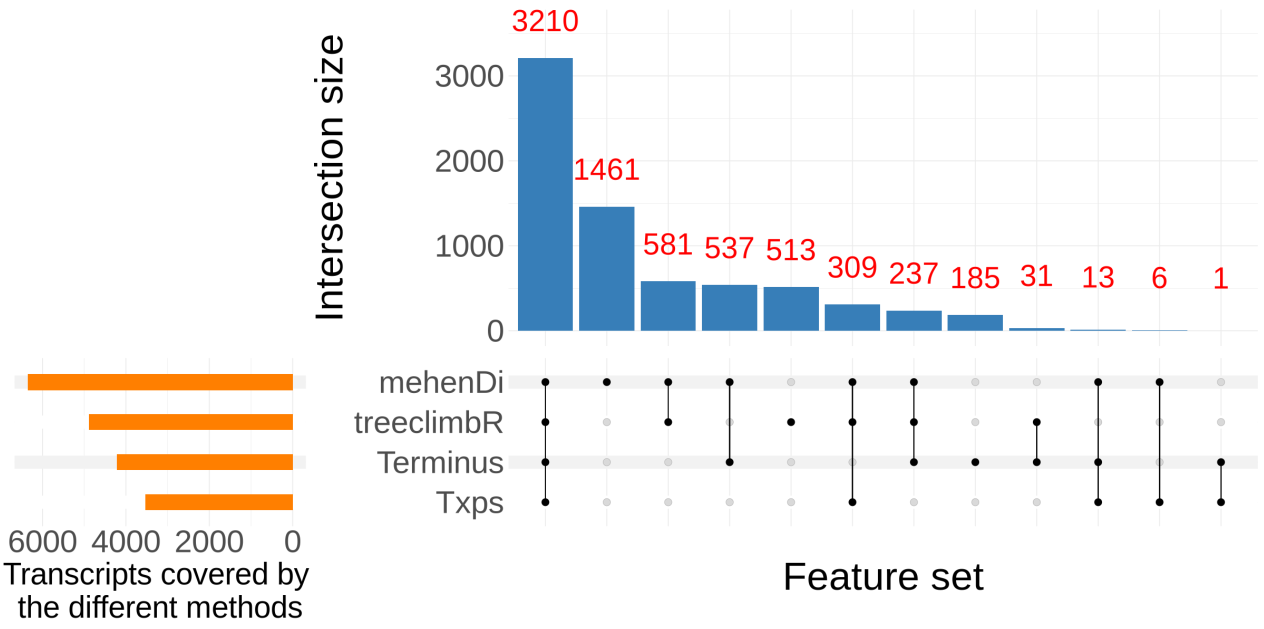

Figure S20: Examining the transcript profile for the gene *CABIN1* in the `ChimpBrain` dataset. A) Transcripts in a pileup style. B) Tree representing the transcripts covered by the gene *CABIN1*, with the red node representing the transcripts covered by the `mehenDi` selected node. C) Inferential replicates for the transcript ENSMUST00000103768, which had the lowest p-value among all the transcripts in the tree. D) Inferential replicates for the `mehenDi` selected node.

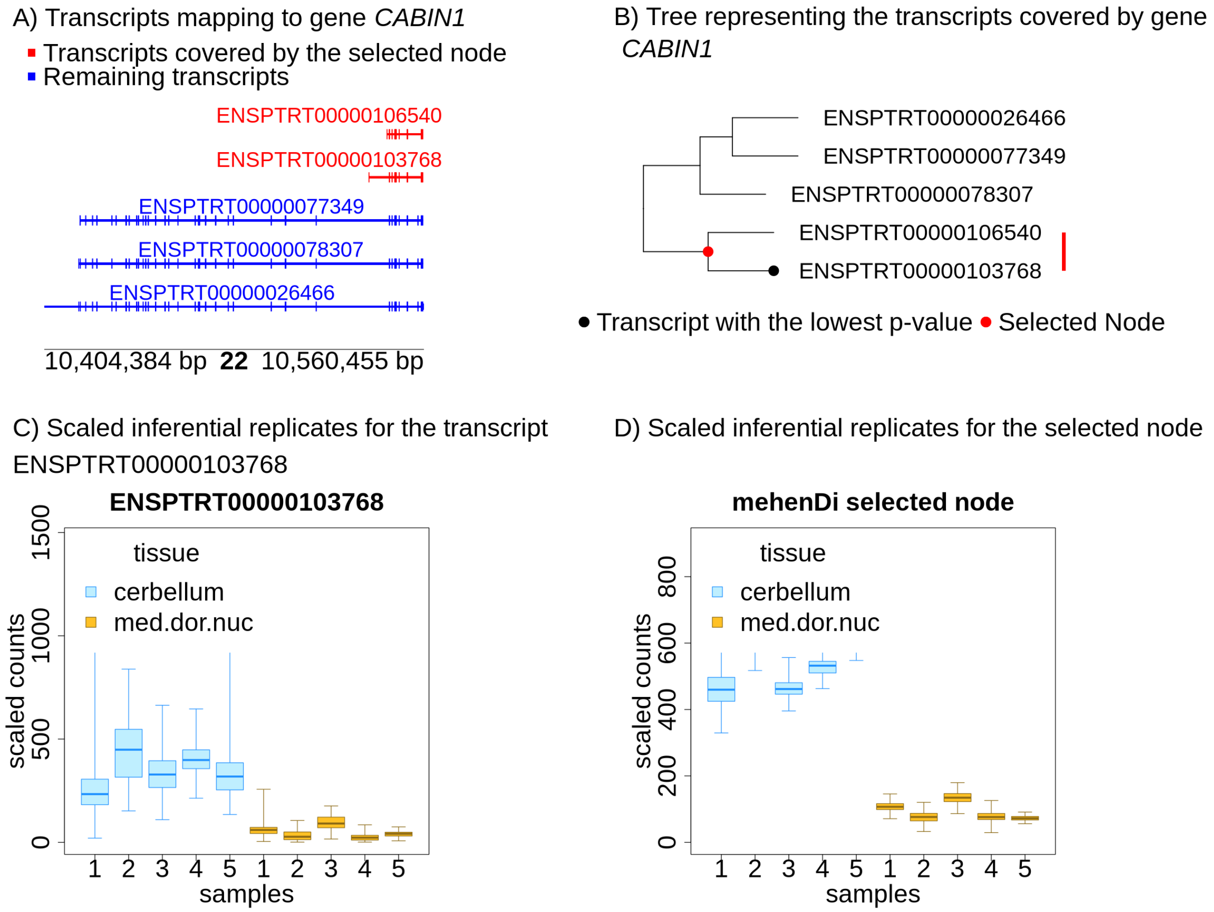

Figure S21: Examining the transcript profile for the gene *CATSPERG* in the `ChimpBrain` dataset. A) Transcripts in a pileup style. B) Tree representing the transcripts covered by the gene *CATSPERG*, with the red node representing the transcripts covered by the `mehenDi` selected node. C) Inferential replicates for the transcript ENSMUST00000020240, which had the lowest p-value among all the transcripts in the tree. D) Inferential replicates for the `mehenDi` selected node.

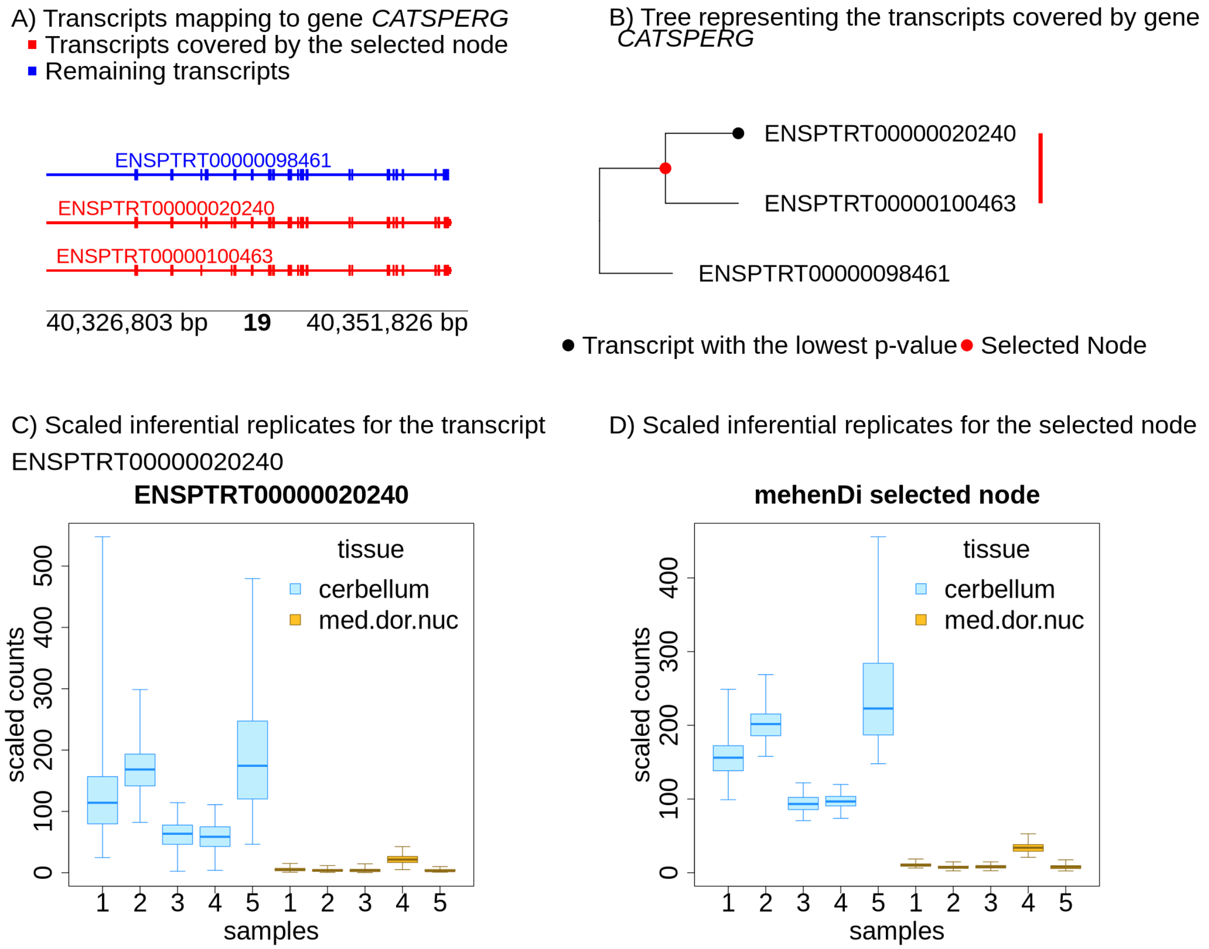

Figure S22: Examining the transcript profile for the gene *EYA1* in the `ChimpBrain` dataset. A) Transcripts in a pileup style. B) Tree representing the transcripts covered by the gene *EYA1*, with the red node representing the transcripts covered by the `mehenDi` selected node. C) Inferential replicates for the transcript ENSMUST00000097751, which had the lowest p-value among all the transcripts in the tree. D) Inferential replicates for the `mehenDi` selected node.

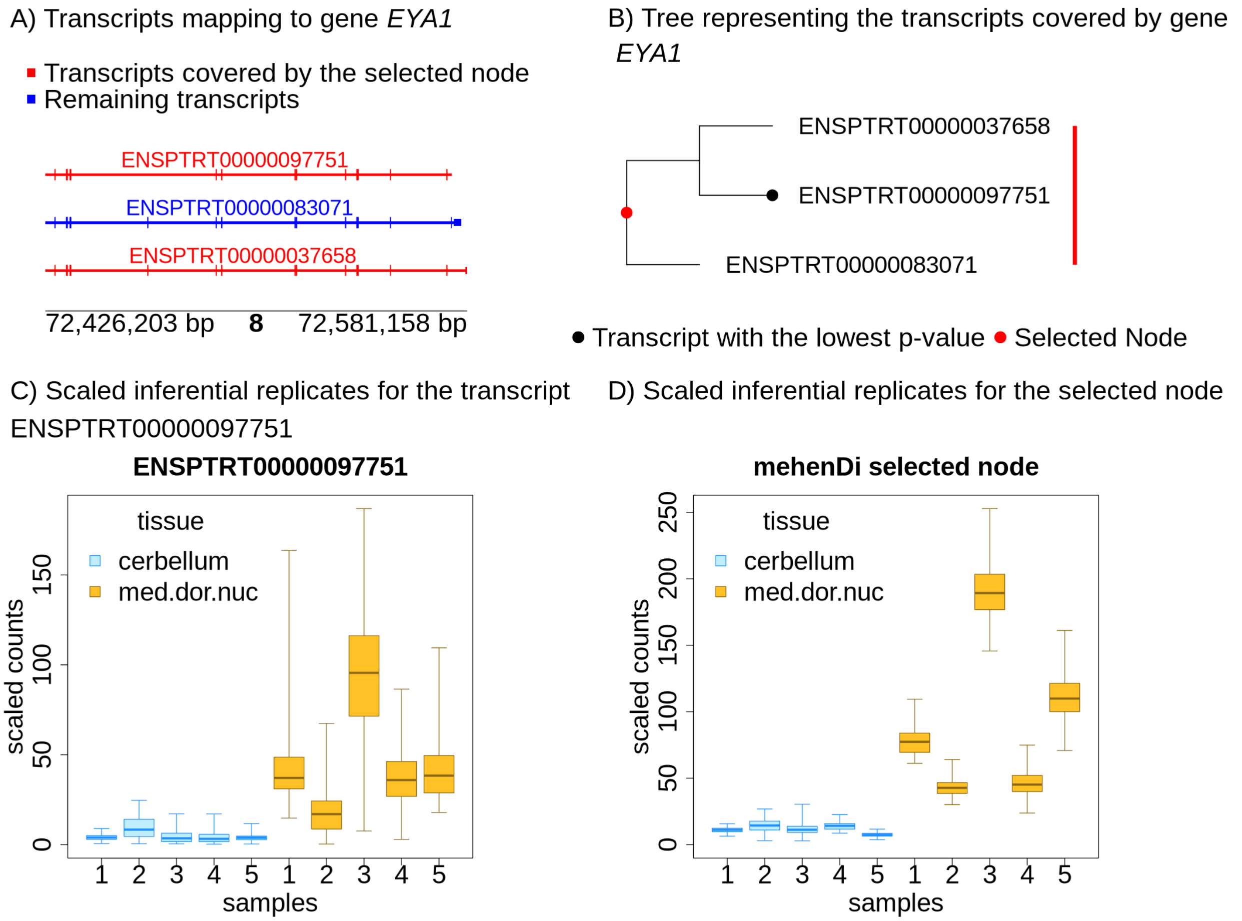

Figure S23: Examining the transcript profile for the gene *MCF2* in the `ChimpBrain` dataset. A) Transcripts in a pileup style. B) Tree representing the transcripts covered by the gene *MCF2*, with the red node representing the transcripts covered by the `mehenDi` selected node. C) Inferential replicates for the transcript ENSMUST00000105901, which had the lowest p-value among all the transcripts in the tree. D) Inferential replicates for the `mehenDi` selected node.

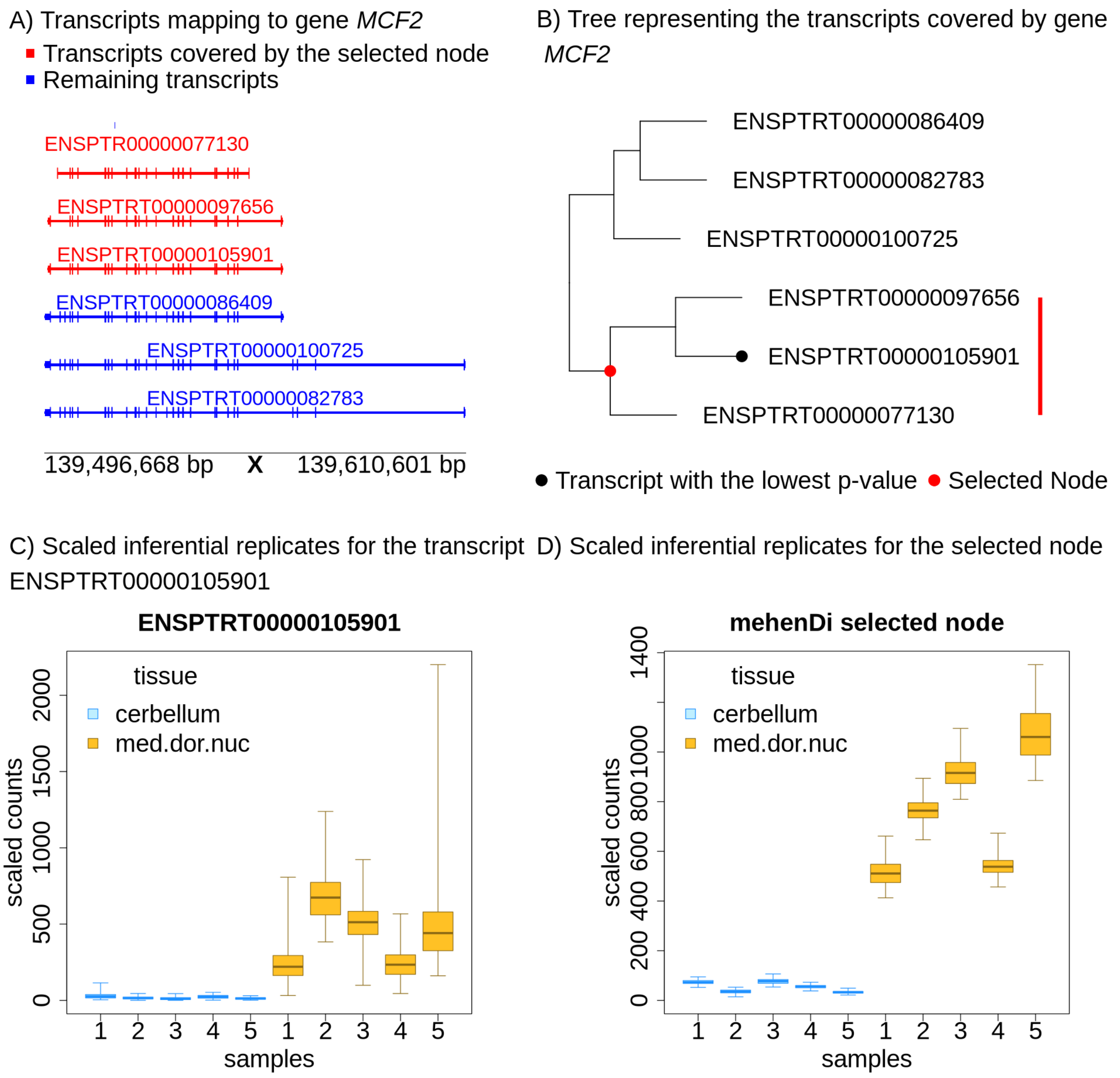

Figure S24: Examining the transcript profile for the gene *MYO5C* in the `ChimpBrain` dataset. A) Transcripts in a pileup style. B) Tree representing the transcripts covered by the gene *MYO5C*, with the red node representing the transcripts covered by the `mehenDi` selected node. C) Inferential replicates for the transcript ENSMUST00000013074, which had the lowest p-value among all the transcripts in the tree. D) Inferential replicates for the `mehenDi` selected node.

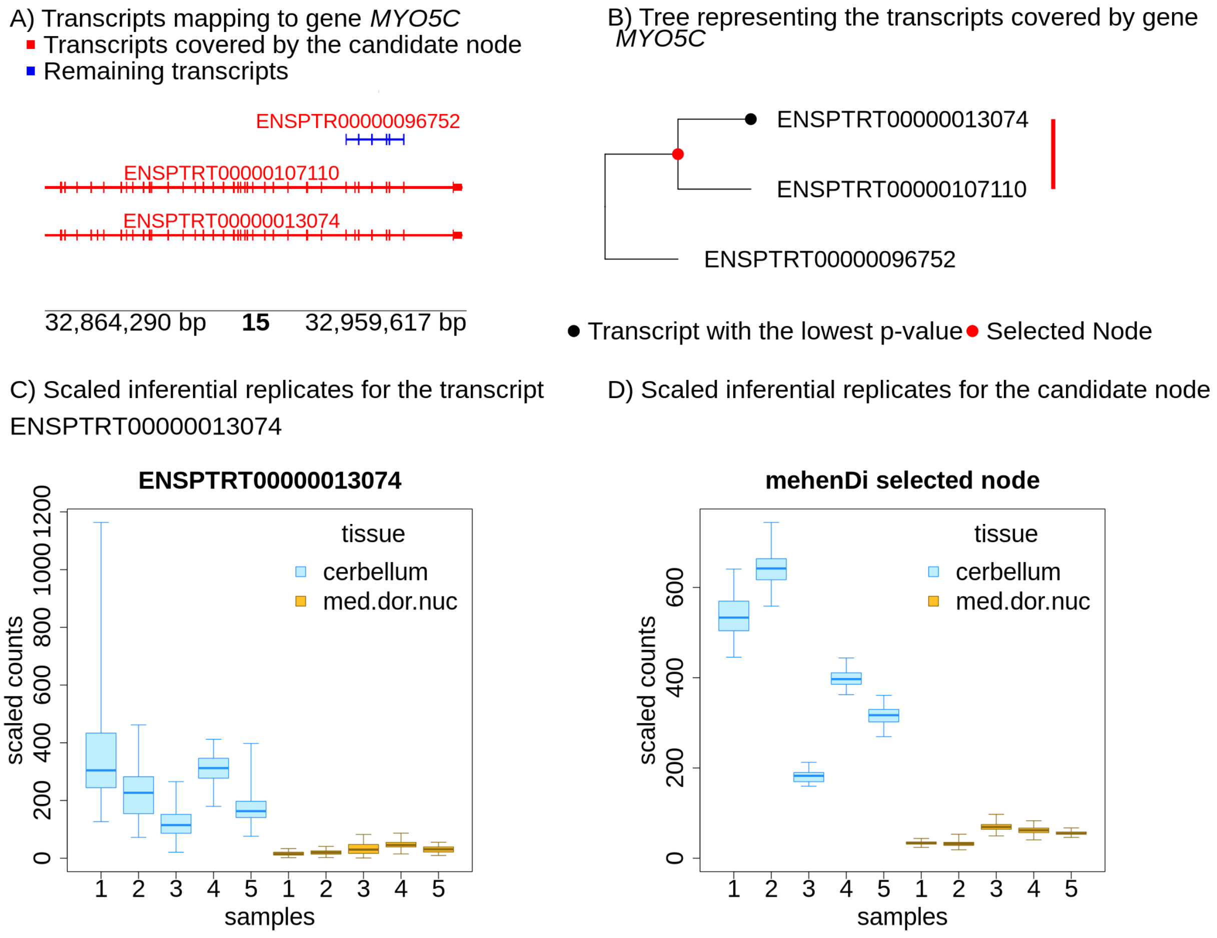

Figure S25: Examining the transcript profile for the gene *PIEZO2* in the ChimpBrain dataset. A) Transcripts in a pileup style. B) Tree representing the transcripts covered by the gene *PIEZO2*, with the red node representing the transcripts covered by the mehenDi selected node. C) Inferential replicates for the transcript ENSMUST00000094046, which had the lowest p-value among all the transcripts in the tree. D) Inferential replicates for the mehenDi selected node.

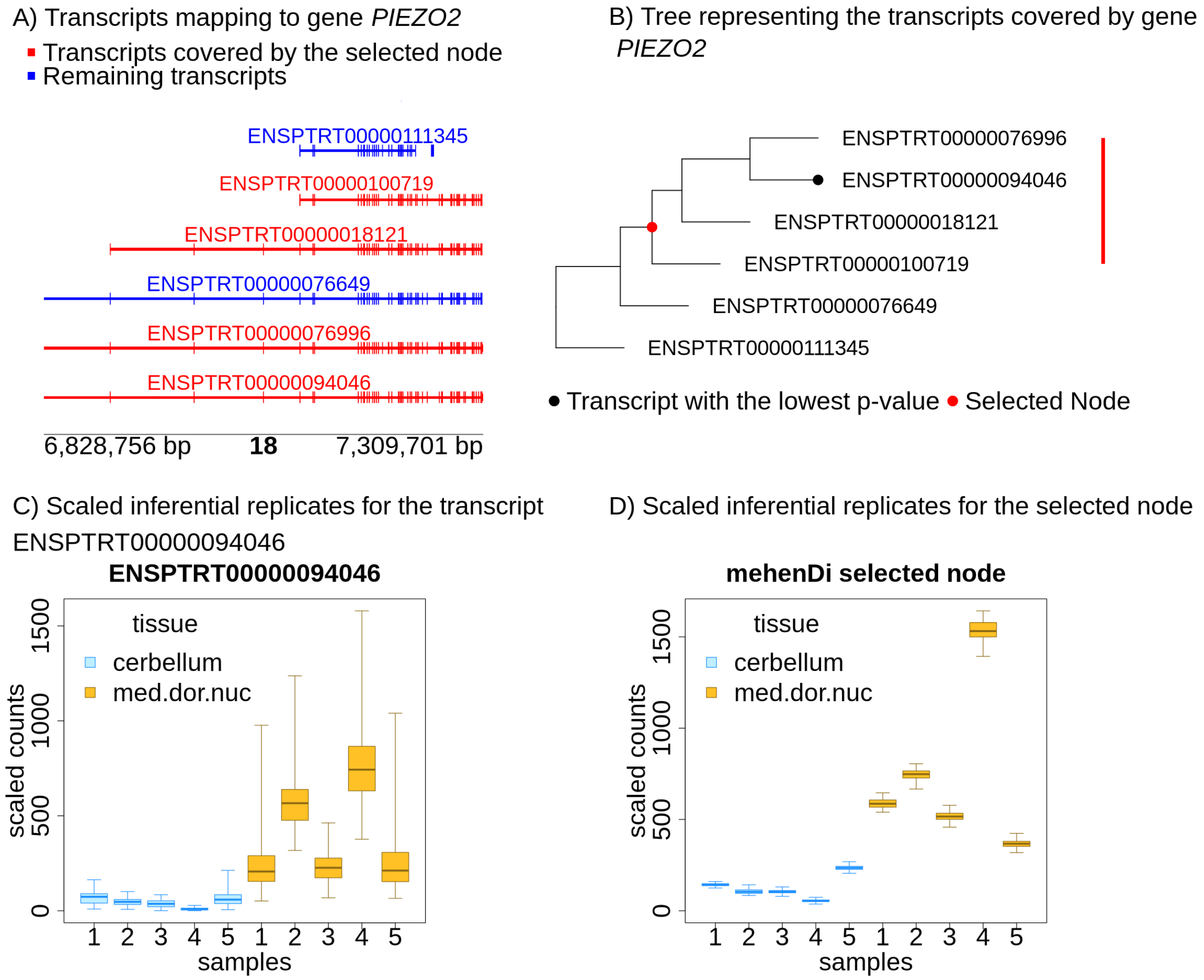
